# Enhancing interpretability of cryo-EM maps with hybrid attention Transformers

**DOI:** 10.64898/2026.01.26.701621

**Authors:** Jie Lin, Ziying Zhang, Yu Zhang, Chengyuan Wang, Guijun Zhang, Xiaogen Zhou

## Abstract

Cryo-electron microscopy (cryo-EM) has become a central technique for determining the structures of macromolecular complexes. However, experimental cryo-EM density maps often exhibit limited interpretability due to noise and heterogeneous, anisotropic resolution, complicating downstream atomic model construction. Here we present DEMO-EMReF, a Swin Transformer-based framework specifically designed for cryo-EM density map refinement that integrates channel attention with hybrid spatial attention. By combining local window-based attention with grid-based sparse global attention, DEMO-EMReF captures fine-grained local density features while modeling long-range, cross-region spatial dependencies, enabling more robust density refinement in heterogeneous and anisotropic map regions. DEMO-EMReF was systematically evaluated on large benchmark sets of both primary maps and half-maps spanning resolutions of 3.0-8.0 Å and was compared against multiple state-of-the-art map enhancement methods. DEMO-EMReF consistently improves resolution-related metrics, map-model correlations, and local atomic resolvability, with particularly robust gains in challenging heterogeneous or anisotropic cases. Importantly, the refined maps facilitate more accurate and efficient automated atomic model building. Together, DEMO-EMReF provides a robust approach for enhancing cryo-EM density maps and enabling more reliable downstream studies.

## Introduction

Recent breakthroughs in single-particle cryo-electron microscopy (cryo-EM) have enabled near-atomic resolution reconstructions of large biomolecular complexes, profoundly transforming the field of structural biology^1,2^. A central objective of cryo-EM is the construction of high-precision atomic models of macromolecular assemblies, a process that critically depends on the quality of the reconstructed three-dimensional density maps^3,4^. However, several intrinsic and technical limitations, including electron dose restrictions, radiation damage, and the inherent flexibility and conformational heterogeneity of macromolecules often lead to reconstructed cryo-EM density maps that suffer from low contrast, high noise, and blurred structural regions^5^. These deficiencies significantly hinder map interpretability and compromise the accuracy of atomic model building.

To improve the interpretability of cryo-EM density maps, a variety of post-processing methods have been developed. Conventional B-factor-based sharpening approaches, implemented in tools such as phenix.auto_sharpen^6^, RELION^7^, and CryoSPARC^8^, enhance high-frequency signals globally but do not account for spatial variations in signal-to-noise ratio, often leading to over-sharpening or under-enhancement in different regions. Localized refinement methods, including LocScale^9^, LocalDeblur^10^, LocSpiral^11^, and OPUS-SSRI^12^, address this limitation by introducing region-specific processing strategies, but typically rely on prior information such as atomic models, masks, or local resolution estimates, which limits their general applicability.

More recently, deep learning-based approaches have been introduced to overcome these constraints. DeepEMhancer^13^ employs a U-Net architecture trained on LocScale-enhanced maps to mimic sharpening effects, whereas EM-GAN^14^ adopts a 3D generative adversarial network trained on simulated density maps derived from atomic models to reduce noise and preserve structural features. EMReady^15^ further introduced a Swin-Conv-UNet (SCUNet)^16^ architecture that integrates residual convolutional layers with Swin Transformer modules and achieves robust improvements in map interpretability, while CryoTEN^17^ employs a Transformer-based UNETR++ architecture^18^ for faster inference. Nevertheless, the reliance of U-Net-based architectures on local convolutional receptive fields makes it challenging to effectively capture long-range dependencies and global structural context in large and heterogeneous maps.

We present DEMO-EMReF, a 3D hybrid attention Transformer architecture that enables long-range information exchange and global context modeling in cryo-EM maps. DEMO-EMReF integrates a channel attention mechanism for global feature recalibration with a hybrid spatial attention module that combines window-based self-attention and a grid-based sparse attention mechanism to overcome the locality constraints of existing methods. This architecture enables selective enhancement of structurally meaningful regions, effective noise suppression, and full-resolution training without information loss induced by downsampling, while also modeling distant structural similarities commonly observed in symmetric or repetitive macromolecular assemblies. Evaluation on diverse test sets of raw experimental primary maps and unprocessed half-maps demonstrates that DEMO-EMReF consistently improves map quality over state-of-the-art methods. Furthermore, atomic model building confirms that structures reconstructed from DEMO-EMReF-enhanced maps exhibit higher accuracy and structural completeness than those derived from deposited experimental maps, highlighting the practical utility of the method for high-precision cryo-EM structural analysis workflows.

## Results

### Overview of DEMO-EMReF

**Figure 1** illustrates the overall architecture and workflow of DEMO-EMReF. The central panel shows a hybrid attention Transformer network specifically designed for cryo-EM density map enhancement. The model consists of four stacked Transformer layers, each containing two key modules. The first is a convolutional Squeeze-and-Excitation^19^ (SE)-based channel attention block that adaptively adjusts channel-wise feature responses to capture global contextual dependencies. The second is a hybrid spatial attention module that integrates local window self-attention including window-based multi-head self-attention (W-MSA) and shifted-window self-attention (SW-MSA) with a grid-based multi-head self-attention (Grid-MSA)^20^ to facilitate long-range interactions across spatially distant regions. This architectural design enables simultaneous modeling of fine-grained local features and long-range correlations in cryo-EM density maps. During training (**Figure 1a**), experimental density maps were collected from the Electron Microscopy Data Bank (EMDB)^21^ and their corresponding atomic models were obtained from the Protein Data Bank (PDB)^22^, covering a resolution range of 3.0-8.0 Å. All maps were preprocessed by normalizing density values and resampling voxel spacing to 1.0 Å. Each map was then partitioned into overlapping density chunks of 64×64×64 voxels, from which 48×48×48 voxel sub-volumes were randomly cropped. The corresponding noise-free simulated density maps generated from the atomic models were processed in the same manner and used as ground-truth targets. The model was optimized using a combined loss function consisting of mean squared error (MSE) and structural similarity index (SSIM), and data augmentation strategies, including random rotations and cropping, were applied to improve the generalization capability of the model. During inference (**Figure 1b**), the target map was divided into multiple overlapping density chunks that were independently processed by the trained model. The enhanced chunks were subsequently stitched together through weighted averaging in the overlapping regions to reconstruct the final enhanced full map. Further details are provided in the Methods section.

**Figure 1.**
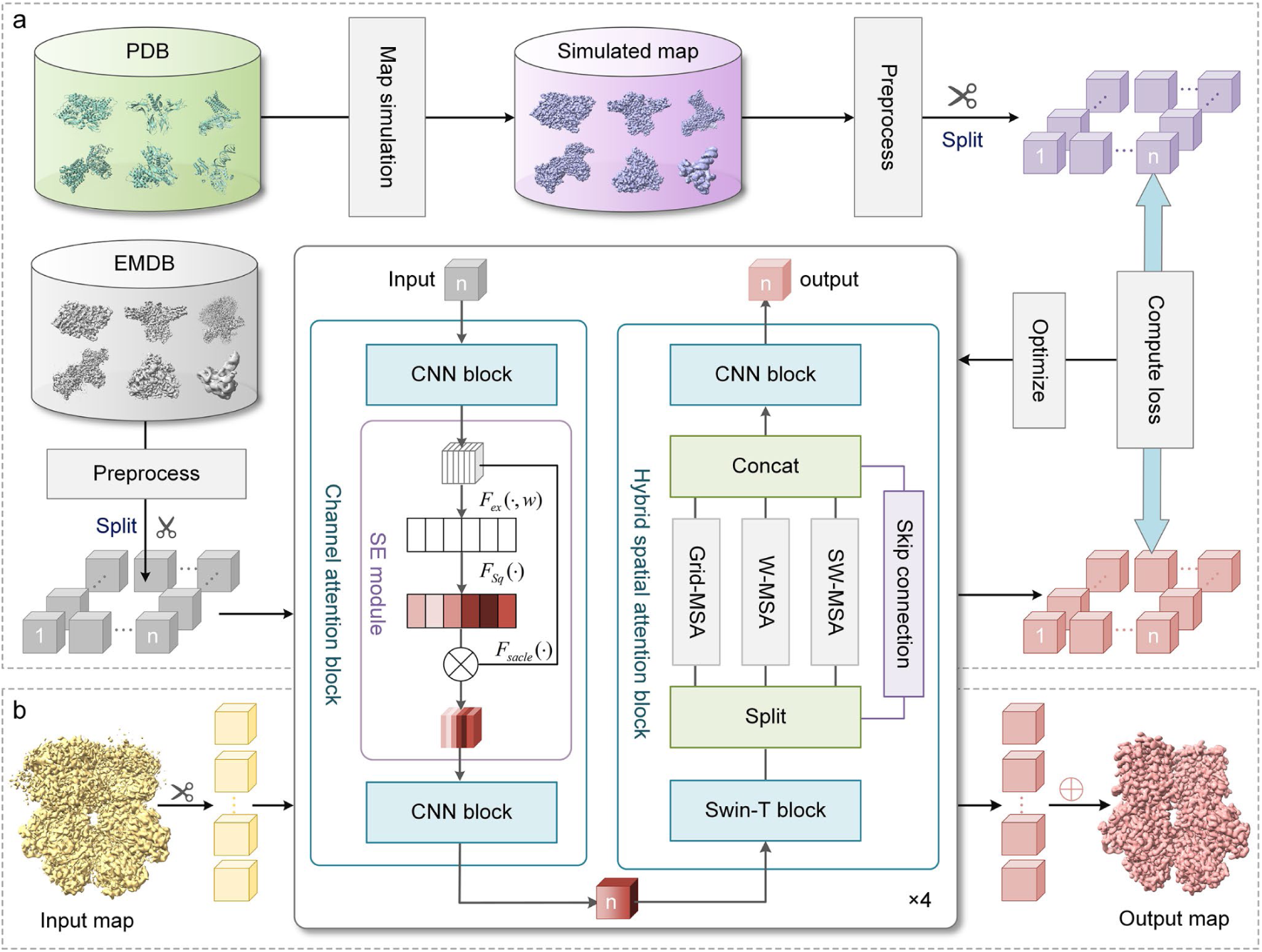
| Flowchart of DEMO-EMReF. **(a)** Training pipeline. Experimental density maps and their corresponding atomic structures were collected from EMDB and PDB, respectively. Noise-free simulated density maps were generated from atomic structures and paired with experimental maps. Both types of maps were preprocessed and partitioned into matched 3D density chunks for supervised training. Each chunk is individually processed by the DEMO-EMReF network (central panel), which integrates convolutional SE blocks for channel-wise attention with a hybrid spatial attention module combining W-MSA, SW-MSA and Grid-MSA. The output chunk is compared with the corresponding simulated chunk to compute the training loss, which is used to update the model parameters via backpropagation. (**b)** Inference pipeline. A target map is divided into chunks, which are processed independently by the trained model. The enhanced chunks are reassembled into the final improved map.

### Improvement of primary maps

The performance of DEMO-EMReF was first evaluated on a benchmark dataset comprising 111 deposited experimental density maps (primary maps) with resolutions ranging from 3.0 to 8.0 Å. For each map, the unmasked map-to-model FSC-0.5 resolution was computed using phenix.mtriage^23^. **Figure 2** and **Supplementary Table 1** summarize the results of the deposited maps before and after refinement by different methods. DeepEMhancer successfully processed 101 out of 111 maps, whereas CryoTEN failed to produce valid unmasked FSC-0.5 values for 2 cases, and one deposited map did not yield a valid unmasked FSC-0.5 value. To ensure a fair comparison, average values were calculated only over test cases where all methods produced valid results. Unless otherwise specified, all reported resolutions correspond to the standard unmasked FSC-0.5 values, and the phenix baseline refers specifically to phenix.auto_sharpen. Overall, DEMO-EMReF significantly improved the average unmasked FSC-0.5 resolution from 5.03 Å for deposited maps to 3.52 Å. As shown in **Figure 2a** and **2b**, DEMO-EMReF enhanced FSC-0.5 resolutions of deposited maps in 97.3% of cases, with 83% of maps exhibiting improvements greater than 0.5 Å. Compared with state-of-the-art methods, DEMO-EMReF (3.52 Å) outperformed EMReady (3.88 Å), CryoTEN (4.45 Å), DeepEMhancer (4.67 Å), and phenix (5.03 Å). Moreover, compared with the strongest baseline method, EMReady, DEMO-EMReF achieved superior FSC-0.5 resolutions in 85.5% of cases (**Supplementary Figure 1a**), with the difference being statistically significant as assessed by a two-sided Wilcoxon signed-rank test. Detailed FSC-0.5 resolutions for all test cases are provided in **Supplementary Table 2**.

**Figure 2.**
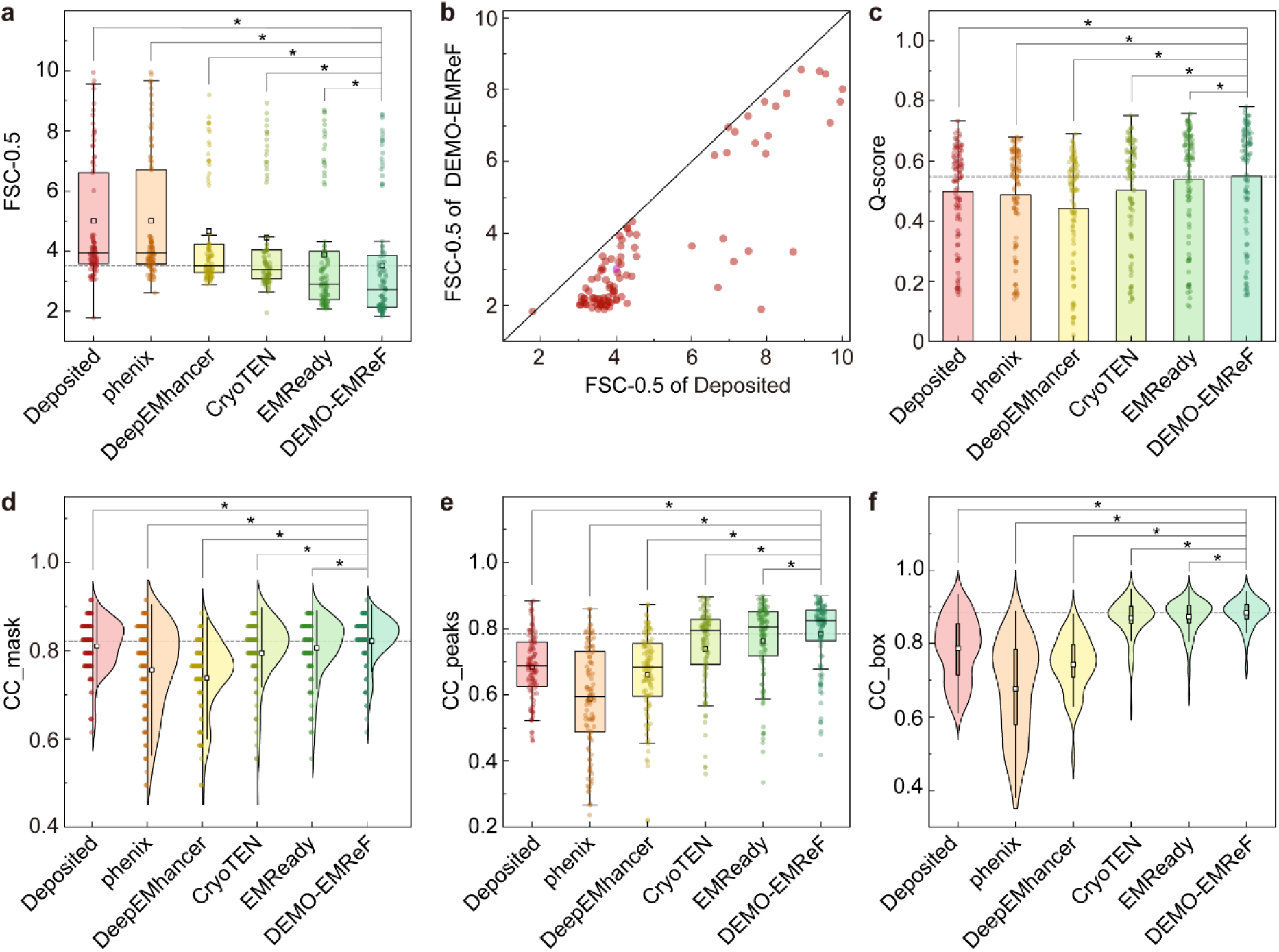
| Comprehensive comparison of map refinement methods on the 111 deposited primary maps. **(a)** Box plots showing the distribution of unmasked map-to-model FSC-0.5 resolutions for each method. The horizontal line within each box denotes the median, square markers indicate the mean, and circles represent individual data points. Asterisks (*) indicate statistically significant differences between DEMO-EMReF and the corresponding method, as determined by a two-sided Wilcoxon signed-rank test (*p* < 0.0001). **(b)** Scatter plot comparing FSC-0.5 resolutions of the deposited maps before refinement and after refinement by DEMO-EMReF. **(c)** Bar chart summarizing average Q-scores for different methods, with individual data points shown as circles. **(d)** Violin plots of CC_mask for each method, overlaid with individual data points and box plots, square markers indicate the mean, and error bars denote the standard deviation. **(e)** Box plots showing CC_peaks values for different refinement methods, with the same statistical annotations as in panel **(a)**. **(f)** Violin plots showing the distribution of CC_box values across different methods.

We further evaluated the interpretability of the enhanced maps using the Q-score^24^ metric implemented in the MapQ plugin of UCSF Chimera^25,26^. The Q-score quantifies the correlation between the density at atomic positions and a reference Gaussian density function and is therefore commonly used to assess local map interpretability at the atomic level. The average Q-score over the entire model has been shown to correlate strongly with map resolution and overall interpretability. The average Q-scores of different methods are summarized in **Supplementary Table 1**. As shown in **Figure 2c** and **Supplementary Figure 1b**, the maps processed by DEMO-EMReF exhibited higher Q-scores than those produced by other methods in most cases, resulting in the highest average Q-score among all methods evaluated. Specifically, DEMO-EMReF improved the Q-score in 99 out of 111 deposited maps **(Supplementary Figure 1c**), yielding an average Q-score of 0.549, which is notably higher than those of the deposited maps (0.498), CryoTEN (0.501), DeepEMhancer (0.442), and phenix (0.486), and marginally higher than EMReady (0.537). Compared to the current state-of-the-art method EMReady, DEMO-EMReF also demonstrated superior Q-scores in 80.2% of cases. Detailed Q-score values for each test case are provided in **Supplementary Table 3**.

To comprehensively evaluate the performance of DEMO-EMReF, we further analyzed the correlation coefficients (CC) between the model-derived and experimental maps, including CC_box, CC_mask, and CC_peaks metrics (**Supplementary Table 1**). As shown in **Figures 2d-f** and **Supplementary Figures 1d-f**, DEMO-EMReF significantly improved all CC metrics for the deposited maps and outperformed other methods. Specifically, the average CC_box, CC_mask, and CC_peaks of the DEMO-EMReF enhanced maps were 0.882, 0.822, and 0.781, respectively, corresponding to improvements of 12.2%, 1.2%, and 14.9% relative to the deposited maps (0.786, 0.810, and 0.680). DeepEMhancer and phenix failed to improve the CC values of the deposited maps on average. CryoTEN achieved average CC_box, CC_mask, and CC_peaks values of 0.868, 0.794, and 0.735, respectively, while EMReady maps obtained corresponding values of 0.872, 0.806, and 0.761. Compared to CryoTEN and EMReady, although the average improvement of DEMO-EMReF is marginal, it is worth noting that correlation-based metrics primarily quantify voxel-wise similarity between maps and can therefore be influenced by shared signal and noise components. Nevertheless, DEMO-EMReF enhanced the CC_box, CC_mask, and CC_peaks in 100, 80, and 98 out of 111 deposited maps (**Supplementary Figures 1g-i**), which are 1.6%, 3.5%, and 6.3% higher than those achieved by CryoTEN and 1.2%, 2.0%, and 2.7% higher than those achieved by EMReady. Detailed CC values for each test case are provided in **Supplementary Table 4-6**.

**Figure 3a** presents the 3.1 Å cryo-EM density map of the Desulfovirgula thermocuniculi IsrB (DtIsrB) complexed with omega RNA and target DNA^27^ (EMD-27533). Although the deposited map is of good quality (FSC-0.5 = 3.79 Å; Q-score = 0.624), portions of the nucleic acid regions display missing or weak density and substantial background noise. DEMO-EMReF markedly improved the map, yielding an FSC-0.5 of 2.23 Å and a Q-score of 0.687. As highlighted in the enlarged view of **Figure 3a**, DEMO-EMReF enhanced the continuity of the DNA phosphate-backbone density and substantially reduced background noise. Accordingly, the processed map obtained significantly higher CC_box/CC_peaks/CC_mask values (0.911/0.859/0.855) than the deposited map (0.688/0.670/0.812). The unmasked map-model FSC curves plotted against inverse resolution (**Figure 3c**) further demonstrate the improvement of the DEMO-EMReF-processed map over the deposited map.

**Figure 3.**
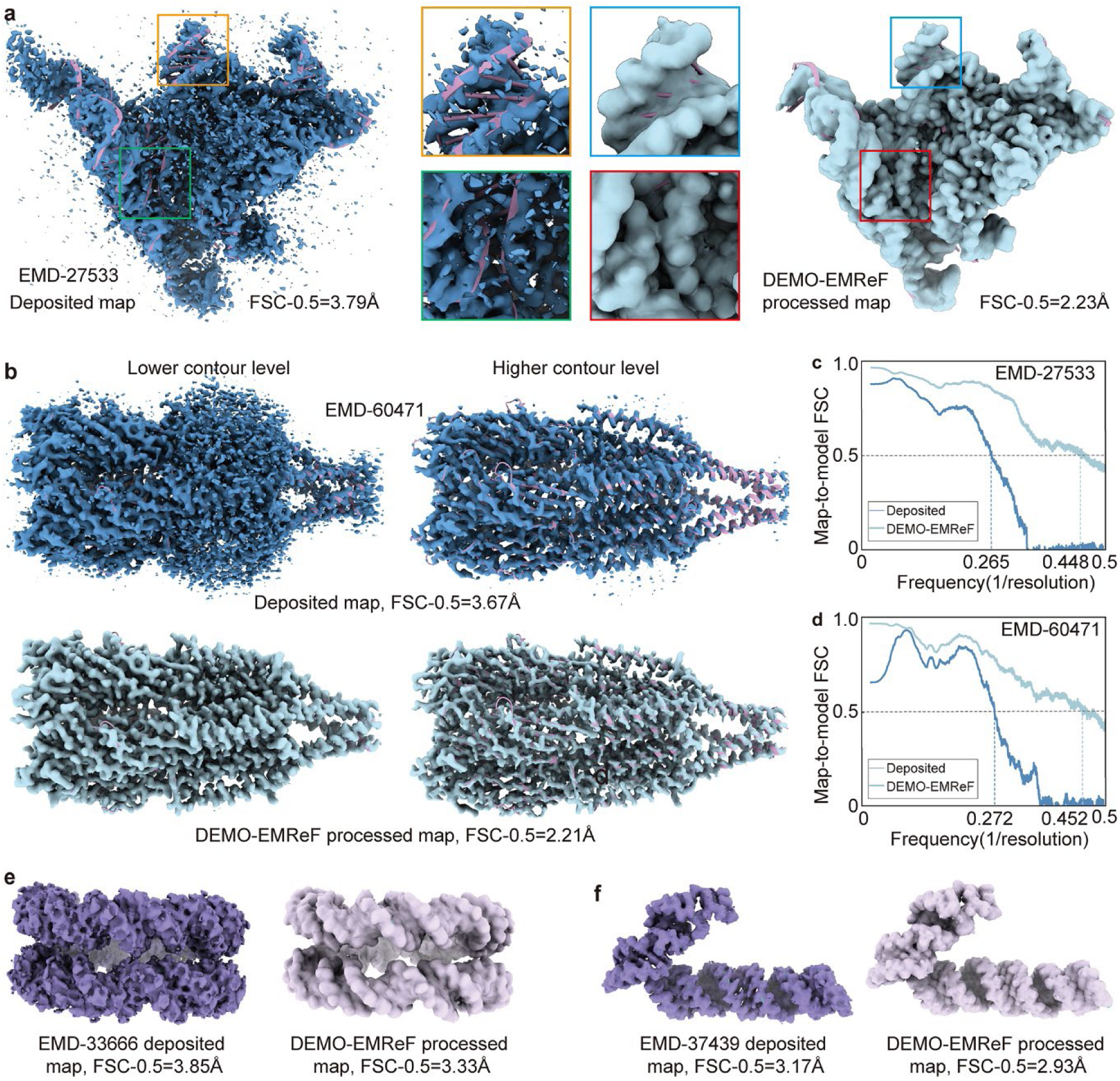
| Examples of density maps enhanced by DEMO-EMReF. (**a**) Cryo-EM density maps of the DtIsrB-omega RNA-target DNA complex (EMD-27533, PDB ID: 8DMB) with a resolution of 3.1 Å before and after DEMO-EMReF enhancement. **(b)** Comparison of the deposited map (EMD-60471, PDB ID: 8ZTS) with a resolution of 3.6 Å and the DEMO-EMReF-enhanced map of the ZAC zinc-activated channel at different contour levels. **(c,d)** Unmasked map–model FSC curves for EMD-27533 and EMD-60471, respectively. **(e)** DNA density in the CENP-A nucleosome-ggKNL2 complex (PDB ID: 7Y7I, chains I and J) from the deposited map(EMD-33666) with a resolution of 3.42 Å and the corresponding enhanced map. **(f)** RNA duplex density of the Cas13h1-crRNA-target RNA complex (PDB ID:8WCE, chains G and T) in the deposited map (EMD-37439) with a resolution of 3.09 Å and enhanced maps.

**Figure 3b** shows a comparison between the deposited map and the DEMO-EMReF enhanced map of EMD-60471, corresponding to the structure of the zinc-activated channel (ZAC) in the Cys-loop receptor superfamily^28^ (PDB ID: 8ZTS), at different contour levels. At the lower contour threshold (left panel), the transmembrane domain of OlZAC is embedded within the surrounding lipid density and is nearly indistinguishable. The DEMO-EMReF enhancement increases the contrast between lipid and protein while suppressing background noise, thereby making the transmembrane domain clearly visible. The right panel shows maps at a higher contour level, highlighting DEMO-EMReF’s ability to enhance high-density features. In the deposited map, one of the four transmembrane helices within the OlZAC subunit appears weakened or even absent, whereas DEMO-EMReF restores these densities, bringing the processed map into close agreement with the PDB model. Quantitatively, DEMO-EMReF improves the unmasked FSC-0.5 resolution from 3.67 Å to 2.21 Å (**Figure 3d**) and raises the Q-score from 0.603 to 0.699. In addition, CC_mask, CC_box, and CC_peaks reach 0.861, 0.901, and 0.857, respectively, which is substantially higher than the deposited map’s 0.791, 0.631, and 0.523.

**Figure 3e** focuses on the DNA structure (chains I and J) from the CENP-A nucleosome in complex with ggKNL2^29^ (PDB ID: 7Y7I, EMD-33666). Although the deposited map exhibits recognizable DNA double-helix features, the overall density boundaries are relatively blurred. DEMO-EMReF substantially reduced background noise, revealing a clearer DNA backbone and more continuous helical features. DEMO-EMReF improved the FSC-0.5 from 3.68 Å to 2.15 Å and enhanced Q-score from 0.442 to 0.498. **Figure 3f** focuses on the RNA duplex region (chains G and T) of Cas13h1 bound to guide and target RNA^30^ (PDB ID: 8WCE, EMD-37439). While the deposited map reveals the overall RNA folding, local density discontinuities are apparent, especially near secondary structure boundaries. DEMO-EMReF markedly enhanced the RNA helical features and density continuity in connecting regions, yielding higher contrast. Accordingly, the DEMO-EMReF-processed map achieved an unmasked FSC-0.5 of 2.09 Å and an average Q-score of 0.674, compared to 3.8 Å and 0.622 for the deposited map. These examples indicate that DEMO-EMReF can effectively enhance key structural features of nucleic acid (e.g., DNA base stacking interactions and RNA secondary structures), facilitating subsequent atomic structure modeling.

### Evaluations on half-maps

Most post-processing methods require half-maps with nominally independent errors in addition to the primary cryo-EM maps. To comprehensively assess the performance of DEMO-EMReF, we further evaluated it on a test set of 27 pairs of half-maps. The average of the two half-maps (averaged map) was used as the input for DEMO-EMReF and the other methods. **Figure 4** and **Supplementary Table 7** report the results of improved maps by different methods. As shown in **Figure 4a**, DEMO-EMReF significantly improved the unmasked map-to-model FSC-0.5 resolution across all 27 half-map cases, outperforming all other methods tested (**Supplementary Figure 2a**), with the average FSC-0.5 improving from 5.02 Å in the averaged map to 3.38 Å after enhancement.

**Figure 4.**
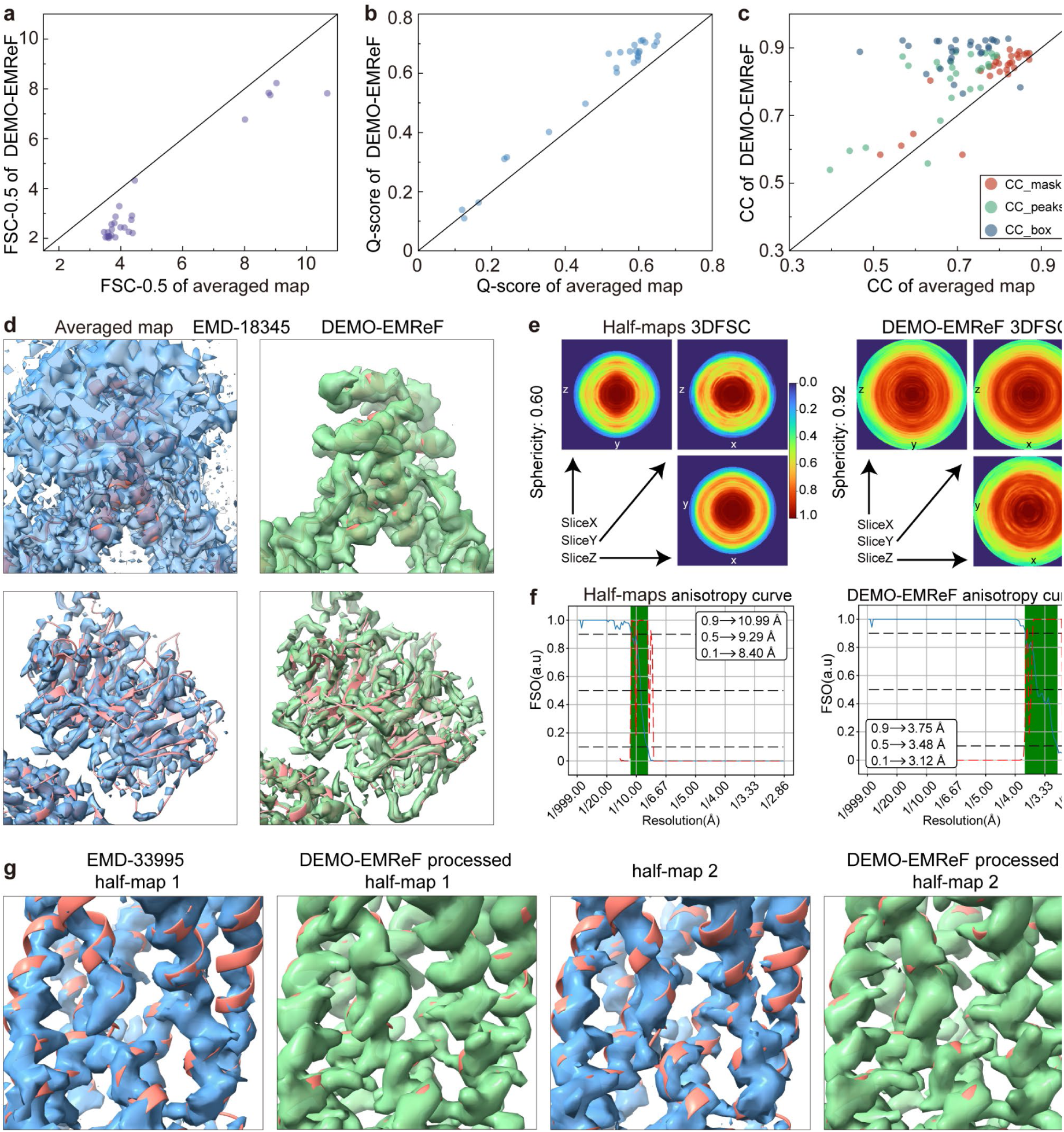
| Results of half-map evaluations. (**a,b,c**) Scatter plots comparing FSC-0.5, Q-scores, and CC values between the original averaged maps and maps refined by DEMO-EMReF. (**d**) Comparison between the original averaged map of EMD-18345 and the DEMO-EMReF-enhanced map at different contour levels. (**e**) Anisotropy analysis of the original half-maps of EMD-18345 and the corresponding DEMO-EMReF-enhanced half-maps. Central slices along the X, Y, and Z directions of the 3DFSC are shown for both the original and enhanced half-maps. (**f**) Anisotropy curves of the original half-maps and DEMO-EMReF-enhanced half-maps, where the FSO (green line) and the *P* value for the Bingham test (red dashed line) were computed using Scipion. a.u., arbitrary units. (**g**) Pairwise comparison of half-maps of EMD-33995, in which one half-map remains unprocessed while the other is enhanced by DEMO-EMReF.

DEMO-EMReF also increased the Q-scores of the averaged maps (**Supplementary Table 7**). As shown in **Supplementary Figure 2b**, DEMO-EMReF obtained an average Q-score of 0.571, surpassing the original averaged maps (0.503) and other post-processing methods (DeepEMhancer: 0.468; phenix: 0.531; CryoTEN: 0.527; EMReady: 0.569). DEMO-EMReF raised the Q-score in 25 out of 27 cases (**Figure 4b**). Further model-to-map correlation analysis (**Supplementary Table 7**; **Figure 4c**) also revealed significant improvement of DEMO-EMReF for the deposited half-map in terms of CC. As shown in **Supplementary Figure 2c**, on average, DEMO-EMReF improved CC_box, CC_mask, and CC_peaks of averaged maps from 0.701 to 0.886, 0.785 to 0.819, and 0.657 to 0.798, respectively. As shown in **Figure 4c**, DEMO-EMReF improved CC_box, CC_mask, and CC_peaks in 26, 25, and 26 out of 27 cases, respectively. Detailed results for each case are provided in **Supplementary Tables 8-12**.

**Figure 4d** presents an example of the DEMO-EMReF-processed averaged map for EMD-18345 (PDB ID: 8QE8). In the original averaged map, only the overall morphology of the complex is visible, with α-helical regions largely buried in background noise (top left). After DEMO-EMReF processing, the map displays well-defined and continuous α-helices. Enlarged views highlight sharply resolved local features with reduced noise and improved contrast relative to the original maps (top right). When both maps are displayed at the same high contour level for the complete map, the processed DEMO-EMReF density retains more density signal along the main-chain tracing path (**Supplementary Figures 2d and 2e**). Quantitative analysis shows an improvement in FSC-0.5 resolution from 8.16 Å to 2.60 Å and an increase in Q-score from 0.621 to 0.678. Consistently, the CC_box, CC_mask, and CC_peaks of the map were improved to 0.897, 0.858, and 0.853, respectively, compared with 0.622, 0.792 and 0.622 for the original averaged map. Additionally, **Figure 4e** shows the three-dimensional Fourier Shell Correlation^31^ (3DFSC) of both the original and processed half-maps of EMD-18345. The original half-maps exhibit strong directional anisotropy with a sphericity of 0.60, whereas DEMO-EMReF processed half-maps increased the sphericity to 0.92, indicating a substantially more isotropic map. Consistently, anisotropy curves based on Fourier Shell Occupancy (FSO)^32^ and Bingham test P-values (**Figure 4f**) show a marked reduction in directional variation, where the resolution gap between the 0.9 and 0.1 cutoff frequencies narrowed, the FSO=1 frequency range broadened, and the FSC resolution at FSO=0.5 improved from 9.29 Å to 3.48 Å. These results demonstrate that DEMO-EMReF effectively suppresses anisotropic artifacts and greatly enhances the overall isotropy and quality of the density map.

To rigorously verify DEMO-EMReF’s ability to extract genuine structural signals from maps, we performed a cross-validation experiment using the 27 pairs of cryo-EM half-maps from the above experiment and 16 additional pairs of cryo-electron tomography (cryo-ET) half-maps. Each half-map was processed independently, and the unmasked FSC-0.5 resolution between the processed half-map and its corresponding unprocessed complementary half-map was computed. As shown in **Supplementary Table 13** and **Supplementary Figure 3a**, DEMO-EMReF improved the average cross FSC-0.5 resolution to 6.18 Å and 6.15 Å for half-map 1 and half-map 2, respectively, compared with 6.86 Å for the unprocessed pairs. A case study on EMD-33995 further confirms that DEMO-EMReF enhances key structural features such as α-helices and β-sheets. The processed half-map shows improved continuity and bridging of previously fragmented densities relative to the deposited half-map (**Figures 4g** and **Supplementary Figures 3b and 3d**). Consistently, the FSC curve between a DEMO-EMReF-processed half-map and the unprocessed complementary half-map exceeds that between the two unprocessed half-maps across the entire resolution range (**Supplementary Figure 3c**), resulting in improved cross FSC-0.5 resolutions of 5.88 Å and 4.55 Å, compared with 7.50 Å for the unprocessed pair. These results further confirm DEMO-EMReF’s ability to enhance true structural signals within half-maps.

### Improved atomic model building

The primary objective of map improvement is to enhance interpretability and thereby improve the accuracy of atomic model building. To evaluate the practical impact of DEMO-EMReF on atomic structure construction, we performed de novo model building on 25 cryo-EM primary maps. Fully automated model building was carried out with phenix.map_to_model^33^ using default parameters, with auto-refinement disabled. The resulting models were compared with deposited PDB structures using phenix.chain_comparison^34^ with a maximum distance threshold of 2 Å. Model quality was evaluated using two metrics: residue coverage and sequence match percentage. As shown in **Figures 5a** and **Supplementary Table 14**, the average residue coverage was improved from 59.4% to 73.6% after DEMO-EMReF processing. Similarly, the sequence match percentage was enhanced from 38.9% to 63.3%. The results demonstrate that DEMO-EMReF substantially enhances map interpretability and enables more accurate atomic model construction. Detailed results are provided in Supplementary Table 15.

**Figure 5.**
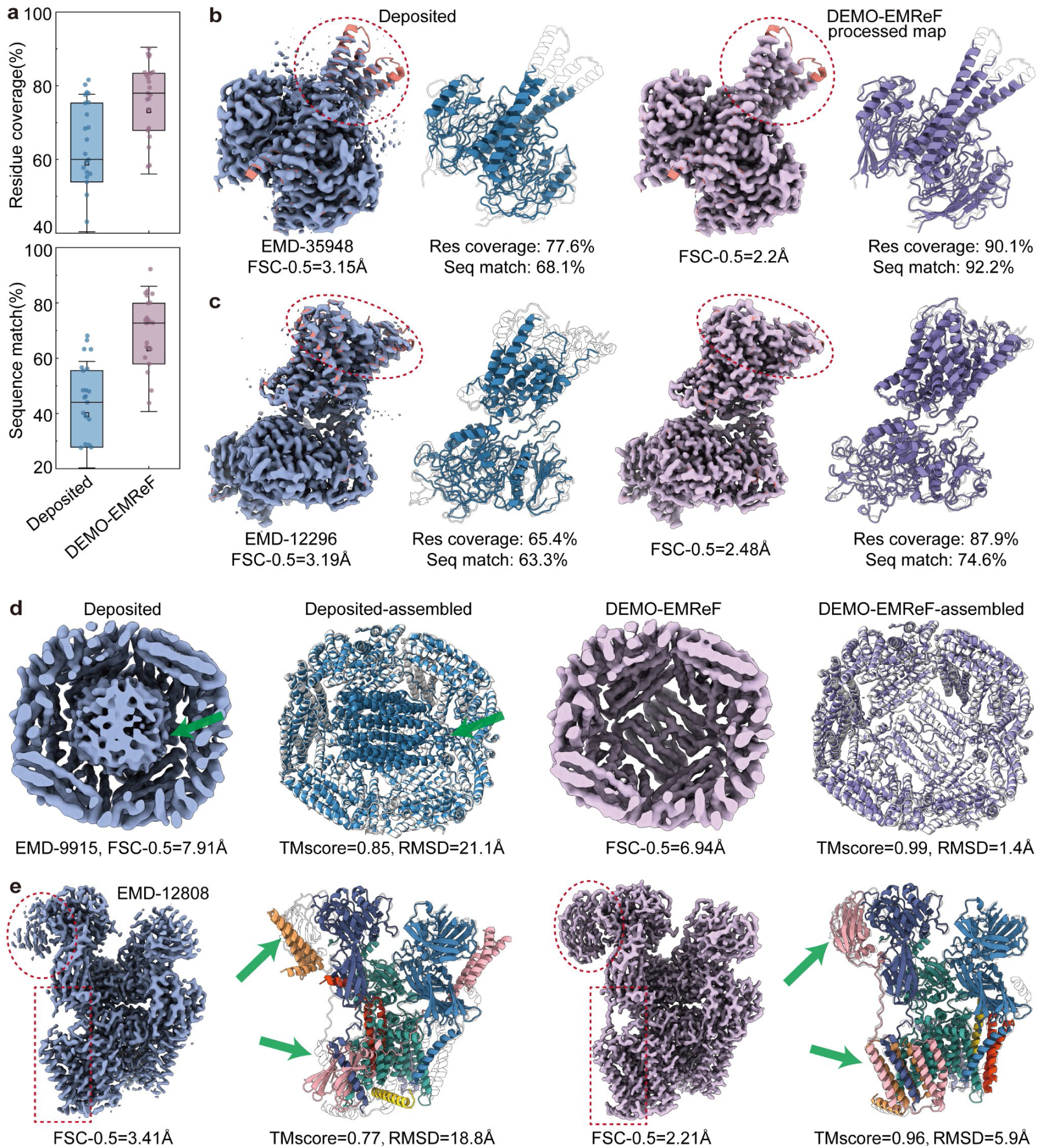
| Improved atomic model building enabled by DEMO-EMReF-enhanced maps. **(a)** Bar plots with overlaid data points comparing residue coverage and sequence match percentage between models built from deposited maps and DEMO-EMReF-enhanced maps. **(b-e)** Comparisons of atomic models built from deposited maps and DEMO-EMReF-enhanced maps for EMD-35948 **(b)**, EMD-12296 **(c)**, EMD-9915 **(d)**, and EMD-12808 **(e)**. Deposited maps and their corresponding model are shown in blue, DEMO-EMReF-enhanced maps and corresponding model in purple, and the reference deposited PDB structure in white. Dashed outlines indicate regions with enhanced density signals after DEMO-EMReF processing. Arrows highlight regions where model assembly was corrected when using DEMO-EMReF-enhanced maps.

**Figures 5b** and **5c** compare atomic models built from the deposited maps with those built from DEMO-EMReF-processed maps for two representative cases. As highlighted by the dashed outlines, the deposited maps exhibit locally weak and discontinuous density, leading to backbone-tracing breaks and ambiguous side-chain placements. In contrast, DEMO-EMReF markedly enhances the signal in these regions, reduces background noise, and improves density continuity, thereby improving backbone tracing and side-chain assignment. **Figure 5b** shows the first example (EMD-35948, PDB ID: 8J2F). The model built from the deposited map achieved 77.6% residue coverage and 68.1% sequence match, whereas the model built from the DEMO-EMReF-processed map reached 90.1% residue coverage and 92.2% sequence match. **Figure 5c** displays another example (EMD-12296, PDB ID: 7NF6). The residue coverage and sequence match of the model generated from the deposited map were 65.4% and 63.3%, respectively, while these metrics were improved to 87.9% and 74.6% when using the DEMO-EMReF-processed map. These results demonstrate that DEMO-EMReF effectively enhances weak density features and overall map continuity, leading to higher-accuracy atomic models in automated de novo model building.

We further assessed the benefits of DEMO-EMReF enhanced maps for structure-based model assembly using two representative cases. Monomeric structures were predicted by AlphaFold3^35^ and assembled into complex models using our previously developed DEMO-EMol^36,37^. The first case is *Bacterioferritin from Streptomyces coelicolor* (EMD-9915, PDB ID: 6K4M), whose deposited map contains low-resolution density around the iron ion (dashed circles in **Figure 5d**). Assembly based on the deposited map resulted in several chains misfitting around the iron-binding region, yielding a complex model with a TM-score^38^ of 0.85 and an RMSD of 21.1 Å. In contrast, DEMO-EMReF effectively filters out spurious density signals related to the iron ions, resulting in a near-native model with TM-score = 0.99 and RMSD = 1.4 Å. The second case is the *yeast Ost6p-containing oligosaccharyltransferase complex* (EMD-12808, PDB ID: 7OCI), whose deposited map exhibits substantial missing structural density (dashed box in **Figure 5e**). Assembly using this raw map produced a complex model with a TM-score of 0.77 and an RMSD of 18.8 Å due to four misfitting chains. When using the DEMO-EMReF enhanced map, the complex was reconstructed with markedly improved accuracy (TM-score = 0.96, RMSD = 5.9 Å), owing to the strengthening of previously ambiguous structural features. In addition, modeling efficiency was markedly improved after DEMO-EMReF processing, as the removal of irrelevant density signals substantially reduced ambiguity during chain placement.

### Assessing density-value optimization

DEMO-EMReF improves map quality by directly optimizing density values in real space. It is therefore important to evaluate the validity of this density optimization. However, direct evaluation is difficult as absolute ground-truth maps or models are typically unavailable in practical applications. Here, we first assessed DEMO-EMReF’s ability to suppress artificially added noise using a benchmark set of primary maps. Specifically, noise-free simulated density maps were generated from the corresponding PDB structures with a grid spacing of 1.0 Å. After normalizing density values to the range of 0.0 to 1.0, Gaussian white noise with a standard deviation of 0.05 was added using Xmipp^39^ to generate noisy maps. These noisy maps were then processed by DEMO-EMReF. The results are summarized in **Supplementary Table 16**, where values closer to those of the simulated maps indicate better density optimization. As shown in the table, all quality metrics of the simulated maps declined substantially after noise addition: CC_box decreased from 0.885 to 0.749, CC_mask from 0.828 to 0.781, and CC_peaks from 0.834 to 0.689. After DEMO-EMReF processing, map quality was largely restored, with CC_box, CC_mask, and CC_peaks improving to 0.890, 0.811, and 0.814, respectively. These values closely approach those of the original noise-free simulated maps (**Supplementary Figure 4a**), indicating effective noise suppression through direct density optimization.

To further evaluate the performance under extreme noise conditions, we introduced Gaussian noise with a standard deviation of 0.2 into a simulated map generated from a PDB structure (PDBID: 8QJX). As shown in **Supplementary Figure 4b**, DEMO-EMReF effectively distinguished noise from authentic structural signals and successfully recovers the original density distribution. Unmasked FSC analysis (**Supplementary Figure 4c**) shows that the global FSC-0.5 resolution improved markedly from 7.46 Å (between the noisy and simulated maps) to 2.49 Å (between the DEMO-EMReF-processed and simulated maps), with consistent enhancement across a broad resolution range. Furthermore, as observed in previous examples (**Figures 3a**, **3b**), DEMO-EMReF can robustly handle real-world noise by directly optimizing density values.

We then evaluated whether DEMO-EMReF overfits the low-resolution regions caused by heterogeneity or flexibility through a representative example. **Figure 6a** shows the deposited structure solved from EMD-22879 at 3.54 Å resolution, where the B-factor values of the RIFIN domains were notably higher than those in the core region of the complex, indicating high structural flexibility in these regions^40^. As shown in **Figure 6b**, these flexible structural regions exhibited relatively lower local map resolution as evaluated by CryoRes^41^. To assess the reliability of density refinement in DEMO-EMReF, we report the local map-to-model correlation in **Figure 6c**, with the left panel showing the deposited map and the right panel showing the DEMO-EMReF-processed map. Several key observations can be made. First, regions with the lowest local correlation (i.e., highest discrepancy) in the DEMO-EMReF-processed map predominantly localize to the two RIFIN domains, which are characterized by lower resolution. In contrast, regions with higher local correlation corresponded to the more rigid parts of MDB1 Fab, where the local resolution was better. This trend is consistent with the local resolution variations observed in the deposited map. Second, compared with the simulated map, DEMO-EMReF enhanced local correlation in structurally reliable regions of MDB1 Fab at approximately 3-5 Å resolution. However, in flexible regions of the RIFIN domains, where the resolution exceeded 5 Å, the density correlation remained comparable to that of the deposited map. This indicates that DEMO-EMReF effectively refines density in well-resolved regions without overfitting poorly resolved or flexible parts, thereby preserving map interpretability and reliability. Overall, the DEMO-EMReF-processed map demonstrated improvements across multiple quality metrics compared with the deposited map: the FSC-0.5 resolution improved from 4.41 Å to 2.81 Å (**Figure 6f**), the Q-score increased from 0.598 to 0.658, and CC_box, CC_mask, and CC_peaks increased from 0.777, 0.833, and 0.754 to 0.929, 0.879, and 0.886, respectively. These results collectively support the rationality and effectiveness of the density refinement in DEMO-EMReF.

**Figures 6.**
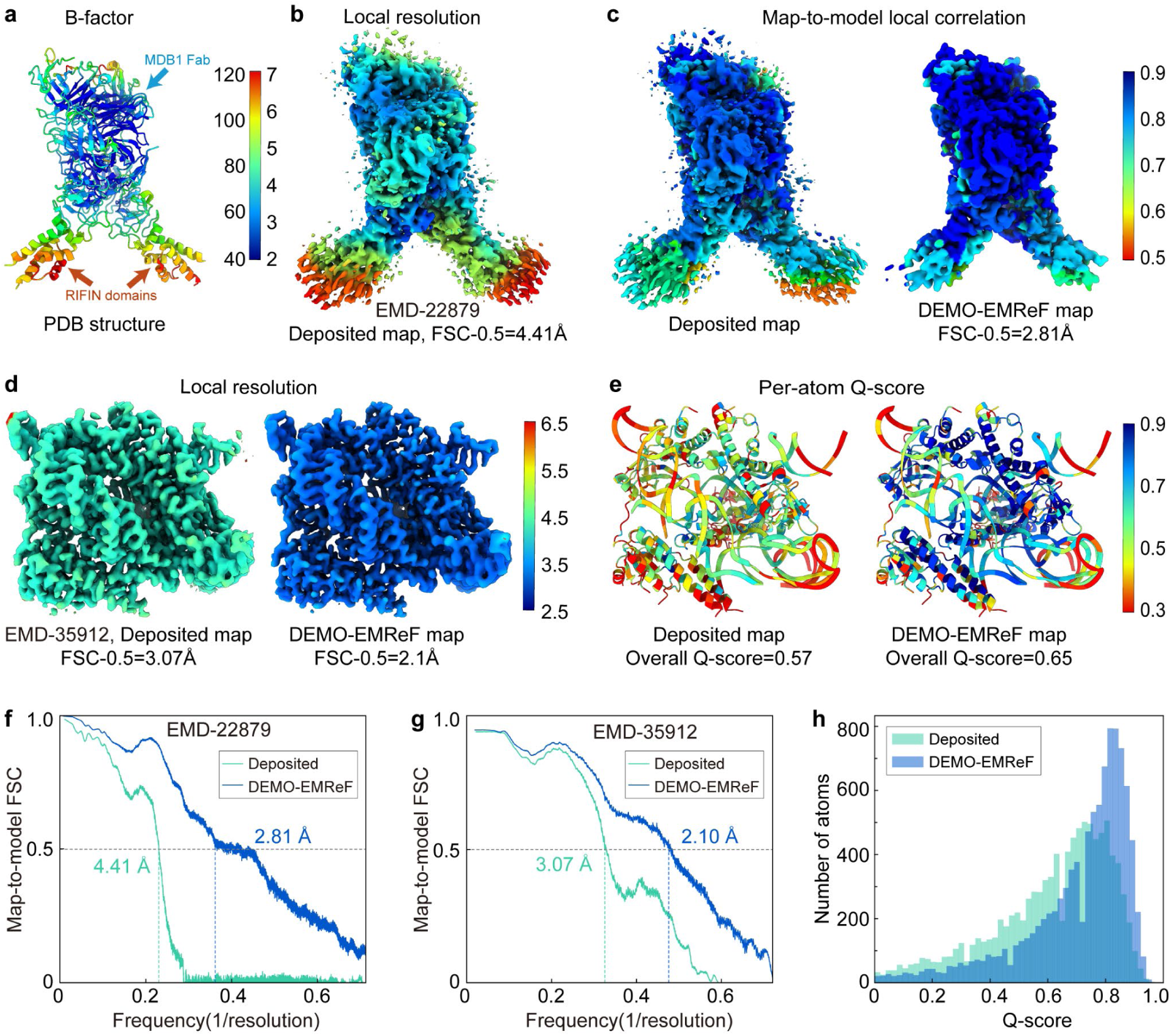
Representative examples illustrating the rationality of density value optimization by DEMO-EMReF. **(a)** Deposited structure (PDB: 7KHF) colored by B-factor. **(b)** Deposited map of EMD-22879 colored by the local resolution. **(c)** Local map-model correlation comparison between the deposited map of EMD-22879 and the corresponding DEMO-EMReF-processed map. **(d)** Local resolution comparison between the deposited map of EMD-35912 and the corresponding DEMO-EMReF-processed map. **(e)** Atomic structures (PDB: 8J12) colored by Q-score computed using the deposited map of EMD-35912 and the DEMO-EMReF-processed map. **(f, g)** Unmasked map-model FSC curves for EMD-22879 and EMD-35912, respectively. **(h)** Q-score distributions of the deposited map (EMD-35912) and the corresponding DEMO-EMReF-processed map.

Finally, we assessed the reasonableness of density optimization for a map with relatively uniform resolution and minimal noise using a representative case. **Figure 6d** (left panel) shows the deposited map of EMD-35912 at 3.08 Å resolution, colored by local resolution. The local resolution of the map is mostly around 3.5 Å, with little apparent noise. The right panel of **Figure 6d** shows the map after DEMO-EMReF processing, where the local resolution is uniformly improved to around 2.5 Å with clearer structural details. The FSC-0.5 resolution of the deposited map was improved from 3.07 Å to 2.10 Å (**Figure 6g**) after DEMO-EMReF processing. **Figures 6e** shows the deposited structures colored by the Q-score calculated from the deposited and DEMO-EMReF-processed maps, respectively, and the corresponding Q-score distributions are shown in **Figure 6h**. The improvement in Q-score is consistent with the enhancement in local resolution. As a result, DEMO-EMReF increased the average Q-score of the deposited map from 0.570 to 0.653. In addition, compared with the CC_box (0.843), CC_mask (0.838), and CC_peaks (0.824) of the deposited map, DEMO-EMReF achieved higher CC_box (0.907), CC_mask (0.863), and CC_peaks (0.866). These results further highlight the reasonableness of density optimization by DEMO-EMReF, demonstrating its ability to improve local map quality without inducing overfitting.

### Ablation experiments

To systematically assess the contributions of key components in DEMO-EMReF, we performed a series of ablation studies. Unlike other deep learning approaches that employ U-Net architectures, DEMO-EMReF adopts an enhanced Swin Transformer that integrates convolution-enhanced SE blocks and sparse grid-window attention to capture complex local and non-local features across both channel and spatial domains. To examine how specific components affect the performance, we trained three ablated variants on the same dataset: (1) a reduced-depth model to quantify the impact of network depth (DEMO-EMReF-3); (2) a model employing SCUNet, which incorporates Swin Transformer blocks into a U-Net backbone; and (3) an SCUNet++-based model^42^ augmented with dense skip connections to strengthen feature propagation.

We evaluated the performance of all ablation models on the benchmark set of primary maps and using FSC-0.5, Q-score, and CC as evaluation metrics. The averaged results of all cases are shown in **Supplementary Table 17**. Although larger input density chunks can enhance a model’s ability to capture non-local features, excessively large chunks drastically increase computational cost. Following recommendations in the EMReady paper, DEMO-EMReF adopts a density chunk of 48×48×48 voxels. To verify the superiority of DEMO-EMReF in capturing non-local features, we compared it against the two SCUNet-based models using the same chunk size. As shown in **Supplementary Figure 5**, the baseline DEMO-EMReF model achieved consistently better performance in terms of average FSC-0.5 resolution and Q-score compared with the SCUNet-based models. This demonstrates that integrating convolution-enhanced SE blocks with sparse grid-window attention enables more stable and effective learning of non-local region features than SCUNet-based architectures.

In general, network depth plays a critical role in determining feature extraction capability. Blindly reducing the number of layers may degrade model accuracy, whereas overly deep networks often require substantially more training data and computational resources. **Supplementary Figure 5** shows that the baseline DEMO-EMReF model, which employs a four-layer network, outperforms the reduced-depth variant DEMO-EMReF-3. Accordingly, we set the network depth of DEMO-EMReF to four layers to balance performance and computational efficiency.

Furthermore, as illustrated in **Supplementary Figure 6** for EMD-27533 (PDB ID: 8DMB), the baseline DEMO-EMReF model effectively reduces the risk of misidentifying genuine density signals near structural boundaries as noise. In contrast, the SCUNet-based model exhibits partial density loss in edge regions (highlighted by dashed outlines), whereas DEMO-EMReF successfully preserves these critical structural features, generating a more continuous and well-defined density distribution. These results further validate the rationality and robustness of the DEMO-EMReF network architecture.

## Discussion

In this study, DEMO-EMReF demonstrates that explicitly facilitating information exchange across distant regions is critical for effective cryo-EM density map refinement. Unlike conventional Swin Transformer architectures that primarily aggregate contextual information through fixed sliding-window attention, the hybrid attention design of DEMO-EMReF enables a more balanced integration of fine local features and long-range structural context. This capability is particularly relevant for cryo-EM map refinement, where accurate modeling of global structural context is critical for mitigating anisotropic resolution artifacts and preserving overall density coherence. In addition, channel-wise feature recalibration allows DEMO-EMReF to adaptively emphasize informative density channels while suppressing noise-dominated responses, thereby improving discrimination between true density signals and background noise. Together, the complementary roles of spatial and channel attention provide an interpretable framework for understanding the improved interpretability of cryo-EM maps observed in our experiments.

Systematic benchmarking on both primary maps and half-maps indicates that DEMO-EMReF consistently improves map quality relative to all compared methods. Beyond enhancing overall resolution and local structural details, the refined maps exhibit reduced anisotropic resolution artifacts, leading to more reliable density interpretation. These improvements translate directly into increased accuracy and efficiency in both de novo atomic model building and fitting-based model assembly. Importantly, ablation analyses suggest that these gains primarily stem from the hybrid attention strategy, in which channel-wise feature recalibration and cross-region spatial interactions jointly contribute to effective modeling of both local and global density features.

Despite these promising results, the performance of DEMO-EMReF could be further improved in several aspects. First, the current training strategy relies on simulated density maps derived from atomic structures as supervision, which may not fully capture the complexity of experimental noise and reconstruction artifacts. Incorporating additional structural information from experimentally determined structures could further enhance model generalization. Second, DEMO-EMReF operates on deposited maps that may have already lost information during reconstruction. Integrating experimental single-particle images, together with simulated images from reference models such as homologous structures or AlphaFold3 predictions, represents a potential avenue for further improvement. Finally, commonly used evaluation metrics such as CC are inherently resolution dependent, and discrepancies in resolution distributions between target and reference maps can bias quantitative assessment. Developing resolution-aware or task-oriented metrics therefore remains an important direction for future work. Overall, given its accuracy and robustness in enhancing both resolution and interpretability, DEMO-EMReF provides a robust and effective tool for improving cryo-EM map quality to support downstream structural and functional studies in the era of cryo-EM structural biology.

## Methods

### Dataset construction

To effectively train and evaluate the DEMO-EMReF framework, we curated a high-quality dataset containing density maps and their corresponding atomic models. Specifically, single-particle cryo-EM and cryo-ET entries with reported resolutions between 3.0 and 8.0 Å were collected from the EMDB. For maps associated with multiple PDB models, only one representative structure was selected to avoid sample redundancy. Entries were excluded if they exhibited non-orthogonal map axes, had mismatches between the structure and the density map, or lacked reported resolution values determined using the FSC-0.143 criterion. To ensure spatial consistency between structures and maps, the phenix.map_model_cc^23^ tool was employed to compute two quality metrics: CC_mask and CC_box. Only samples with CC_mask > 0.75 and CC_box > 0.6 were retained. For maps with reported resolutions of 6.0-8.0 Å, the CC_mask threshold was moderately relaxed to 0.6 to expand the dataset.

In addition, to eliminate redundancy, the FASTA sequences of all samples were clustered using MMseqs2^43^ at 80% sequence identity, and one representative from each cluster was retained. The resulting dataset comprised 923 non-redundant map-model pairs. Among these samples, 111 cryo-EM maps were randomly assigned to the test set, and 16 cryo-ET cases for which half-map files could be successfully retrieved from the EMDB and exhibited good agreement with their atomic models were also included in the test set. The remaining data were divided into the training set and validation set in a 4:1 ratio.

### Map data preprocessing

To ensure consistent preprocessing across all maps and improve training efficiency, a unified preprocessing protocol was applied to both experimental maps and their corresponding simulated maps. All maps were resampled to a uniform voxel spacing of 1.0 Å using cubic interpolation. Negative density values were clipped to zero to remove non-physical noise. To normalize density amplitudes across different maps, all density values were linearly scaled to the range [0, 1] using the 99.999th percentile of each map’s density distribution. In the training set, the input consisted of deposited experimental maps, while the supervision was provided by simulated density maps generated from the corresponding PDB structures. These simulated maps were generated using a Gaussian kernel-based method, in which each atom contributes to the density at a grid point according to its atomic number and spatial distance. Specifically, the density value at a grid point *x* is computed as:

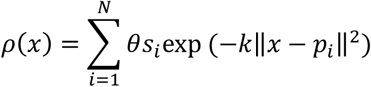

where *s_i_* and *p_i_* denote the atomic number and Cartesian coordinates of the *i-*th atom, respectively, and *N* is the total number of atoms. The attenuation factor *k* was determined from the reported FSC resolution *R*_0_ at the 0.143 threshold^44^ as *k*=(*π*∕0.9*R_0_*)^2^, and the scaling factor was defined as *θ*=(*k*∕*π*)^3⁄2^. In addition, given the large size of cryo-EM maps, all maps were partitioned into overlapping 3D density chunks to enable efficient training and inference. Only chunks containing meaningful density signals were retained, while empty regions were discarded. During validation and testing, maps were split into overlapping chunks of size 48×48×48 with a stride of 24 voxels. The final enhanced maps were reconstructed by weighted averaging in the overlapping regions.

### Neural network architecture

DEMO-EMReF adopts a Transformer-based neural network architecture with a hybrid attention mechanism, designed to enhance both the resolution and resolvability of density maps. As illustrated in **Figure 1b**, the model takes as input a 3D density chunk of size 48×48×48 voxels (voxel spacing of 1.0 Å) and outputs an enhanced density chunk of the identical dimensions. The overall architecture (**Figure 1**, center panel) consists of four sequentially stacked residual hybrid attention modules, each comprising two core components: a channel attention module and a hybrid spatial attention module.

The channel attention module is built upon the SE mechanism, which dynamically modulates feature responses along the channel dimension. Specifically, the input density chunks are first processed by a 3×3×3 convolutional layer with an expansion factor of six to increase channel dimensionality. A global squeeze operation with a reduction ratio of 0.5 is then applied to aggregate global context, followed by a 1×1×1 convolution (excitation) to restore the original channel dimension. Through global context compression and channel-wise reweighting, this module enhances informative features while suppressing less relevant ones, thereby improving feature discrimination and signal-to-noise characteristics.

To capture both local and long-range spatial dependencies in density maps, DEMO-EMReF employs a hybrid spatial attention module built upon the Swin Transformer framework. An initial Swin Transformer block is applied to extract local and non-local spatial representations. To further expand the effective receptive field and strengthen cross-region interactions, feature maps are partitioned and processed using a Grid-MSA together with W-MSA and SW-MSA. In this process, feature maps are divided into non-overlapping grid regions with small spatial offsets, enabling sparse attention computation across grids and facilitating efficient long-range information exchange. The resulting features are subsequently integrated through a multi-layer perceptron and a convolutional fusion module to combine local and global representations.

Overall, the network synergistically integrates channel-wise feature recalibration with an enhanced spatial attention, enabling simultaneous learning of fine-grained local details and long-range structural dependencies. This design allows DEMO-EMReF to robustly refine cryo-EM density maps across a wide range of resolutions and structural complexities.

### Network training

The network of DEMO-EMReF was trained for 100 epochs on an NVIDIA A800 GPU with 80 GB of memory. To balance convergence stability and memory efficiency, a batch size of 8 and a 4-layer Transformer encoder-decoder architecture were adopted. All maps were split into overlapping density blocks of size 48×48×48 voxels after preprocessing, consistent with the block size used during inference. To reduce overfitting and improve generalization, several data augmentation strategies were applied, including random cropping from 64×64×64 to 48×48×48 voxel chunks, random 90° rotations, and axis flipping. In addition, Gaussian white noise with a variance of 0.1 was added to each density block with a probability of 0.1 to enhance the denoising capability of the model. Dropout layers with a rate of 0.1 were applied in both the Transformer encoder and decoder modules.

The training loss function combined the MSE^45^ loss and the SSIM^46^ loss. The MSE loss quantifies voxel-wise local differences and is defined as:

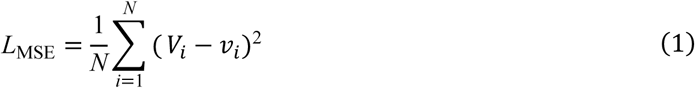

where *N* denotes the total number of nonzero voxels (i.e., voxels that are nonzero in either the predicted or target blocks), *V*_𝑖_ is the predicted voxel value, and *v*_𝑖_ is the corresponding ground-truth value.

To capture global structural similarity, the SSIM loss was incorporated and computed as:

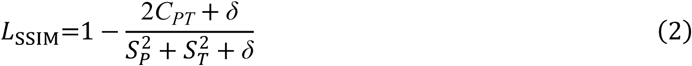

where 𝑆_𝑃_and 𝑆_𝑇_denote the standard deviations of the predicted and target density chunks, respectively, *C_PT_* is their covariance, and *δ* =10^−6^ is a small constant to prevent division by zero. The total loss function is defined as a weighted sum of the two terms:

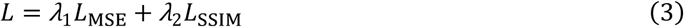

where *λ*_1_ = 1 and *λ*_2_ = 4 are weighting factors used to balance their contributions.

Model optimization was performed using the Adam optimizer with an initial learning rate of 5×10^-4^, which was decayed to a minimum of 1×10^-5^ using a cosine annealing schedule over the course of training. Model performance was evaluated on the validation set after each epoch, and the checkpoint with the lowest validation loss was selected for final evaluation. The training process was implemented using PyTorch 1.8.1 and CUDA 11.1. All hyperparameters were empirically determined based on validation performance and computational efficiency.

### Evaluation metrics

#### Frequency-domain resolution and anisotropy

FSC curves between maps (before and after enhancement) and their corresponding atomic models were calculated using phenix.mtriage^23^, with the resolution at FSC = 0.5 adopted as the primary metric for map resolution. To assess directional resolution anisotropy, 3DFSC was adopted. 3DFSC provides voxel-wise visualization of directional FSC values across Fourier space, enabling detection of anisotropic regions arising from preferred particle orientation or data-collection geometry. The sphericity derived from the 3DFSC distribution was used as a quantitative descriptor of isotropy and directional resolution variation in the enhanced maps.

In addition, FSO was also employed to evaluate frequency-dependent anisotropy and completeness of the enhanced maps. FSO measures the fraction of directions in each Fourier shell for which the directional FSC exceeds 0.143. Values close to 1 indicate an isotropic Fourier signal distribution, whereas lower values reflect increasing anisotropy. The frequency over which FSO declines from 0.9 to 0.1 defines the anisotropy transition zone, and the shell with FSO=0.5 approximately corresponds to the global resolution. Thus, FSO complements FSC and 3DFSC by providing a compact descriptor of both resolution and directional uniformity.

#### Real space map-model correlation

Real-space correlation coefficients (CC_mask, CC_box, and CC_peaks) were computed using phenix.map_model_cc to quantify the agreement between density maps and atomic models. CC_mask measures correlation within a molecular mask, CC_box assesses global correlation across the entire map, and CC_peaks evaluates fit at the highest-density regions by comparing prominent density peaks.

#### Local atomic resolvability

Local model interpretability was evaluated using Q-score, computed with the MapQ plugin in UCSF Chimera. Q-score quantifies the similarity between the observed local density profile around an atom and an idealized Gaussian-shaped peak, with higher values indicating clearer atomic features and higher local resolvability. In addition, local resolution was estimated using CryoRes, a deep-learning method that predicts voxel-wise resolution directly from a map. For each voxel, CryoRes outputs a resolution value learned from FSC-based local-resolution training data, producing a 3D map that reflects regional variations in density quality, where lower values indicate better-resolved features.

#### Accuracy of reconstructed atomic models

The accuracy of de novo reconstructed atomic models from maps was assessed using residue coverage and sequence matching rate calculated by phenix.chain_comparison. Residue coverage quantifies the percentage of residues whose representative atoms (Cα for proteins; P for nucleic acids) lie within 3.0 Å of corresponding residues in the reference structure, regardless of residue identity. Sequence matching rate measures the proportion of residues that are both spatially aligned and correctly assigned to the reference sequence. Higher values for both metrics indicate better map interpretability, more accurate backbone placement, and more reliable residue typing.

For atomic models assembled using computationally predicted chain structures, structural accuracy was evaluated by TM-score and RMSD calculated with US-align^47^. TM-score evaluates the global topological similarity between two protein structures on a scale of (0, 1], with higher values indicating closer overall agreement. RMSD measures the average distance between corresponding atoms after optimal alignment, with lower RMSD values denoting greater structural similarity.

### Performance evaluation of DEMO-EMReF

The performance of DEMO-EMReF was evaluated on a benchmark set of 111 primary maps and compared with DeepEMhancer, EMReady, CryoTEN, and phenix, under identical hardware conditions using default parameters for all methods. Average performance metrics were computed only for cases that yielded valid outputs across all methods. DeepEMhancer provides three model configurations: tightTarget, wideTarget, and highRes. For maps with resolutions between 3 and 6 Å, the best-performing result among the three models was selected, whereas for maps in the 6-8 Å range, only the tightTarget and wideTarget models were considered.

To examine the robustness of DEMO-EMReF under low signal-to-noise conditions and its ability to recover true structural signals, an independent evaluation was performed on 43 half-map pairs (27 EM pairs and 16 ET pairs). DEMO-EMReF was first applied to the averaged map from each half-map pair. In addition, a cross-half-map validation was performed, in which one half-map was processed by DEMO-EMReF and FSC was computed between the processed half-map and its unprocessed complementary half-map.

To evaluate the benefits of DEMO-EMReF for atomic model building, de novo model building was performed on 25 enhanced maps using phenix.map_to_model. The resulting models were assessed using phenix.chain_comparison and compared with models built from the corresponding deposited maps. The effect of DEMO-EMReF on structure-based assembly was further evaluated by assembling complex structures from AlphaFold3-predicted monomers using our previously developed DEMO-EMol pipeline. Models assembled using DEMO-EMReF-enhanced maps were compared with those generated from deposited maps.

To assess the reasonableness of the density optimization performed by DEMO-EMReF, artificial Gaussian noise was added to simulated maps generated from the PDB structures corresponding to the 111 primary maps using Xmipp. DEMO-EMReF was then applied to the noisy maps to evaluate its ability to recover the simulated maps by identifying the noise. Furthermore, DEMO-EMReF was tested on representative cases exhibiting low-resolution regions caused by structural heterogeneity or flexibility, as well as on cases with relatively uniform resolution and minimal noise, to assess its ability to improve local density quality.

Finally, to quantify the contributions of individual architectural components in DEMO-EMReF, ablation studies were conducted under identical experimental settings, using the same datasets and hyperparameters as the baseline DEMO-EMReF model. Three ablated variants were examined: (1) DEMO-EMReF-3, a reduced-depth version of the baseline model; (2) a SCUNet-based model incorporating Swin Transformer blocks into a U-Net backbone; and (3) a SCUNet++-based model enhanced with dense skip connections.

## Data availability

The authors declare that the data supporting the findings and conclusions of this study are available within the paper and its Supplementary Information. The DEMO-EMReF model weights, along with the density maps used for training and testing, are available at https://github.com/zhouxglab/DEMO-EMReF. The experimental density maps and the deposited structures used in this study were downloaded from EMDB and PDB. Additional data are available from the corresponding author upon reasonable request.

## Code availability

The source code and corresponding data are freely available for academic or non-commercial users at https://github.com/zhouxglab/DEMO-EMReF.

## Acknowledgements

This work was supported by the National Key R&D Program of China (2022ZD0115103 to G.Z), the National Nature Science Foundation of China (62203389 to X.Z., 62173304 to G.Z.), Fundamental Research Funds for the Provincial Universities of Zhejiang (RF-C2024006 to X.Z.), Leading Innovative and Entrepreneur Team Introduction Program of Zhejiang (2023R01006 to G.Z.), Zhejiang Provincial Special Support Program for High-Level Talents (2023R5248 to X.Z.), and the “Pioneer” and “Leading Goose” R&D Program of Zhejiang (2025C01121 to X.Z., 2025C01190 to G.Z.).

## Author contributions

X.Z. conceived and designed the project. J.L. developed and implemented the computational pipeline and performed the test. G.Z. and C.W. jointly supervised the study. Z.Z. performed data analysis and prepared the figures. Y.Z. assisted with data analysis. J.L. and X.Z. wrote the manuscript. All authors revised and approved the final manuscript.

