## Supplementary Information for "Enhancing interpretability of cryo-EM maps with hybrid attention Transformers"

### Supplementary Figures

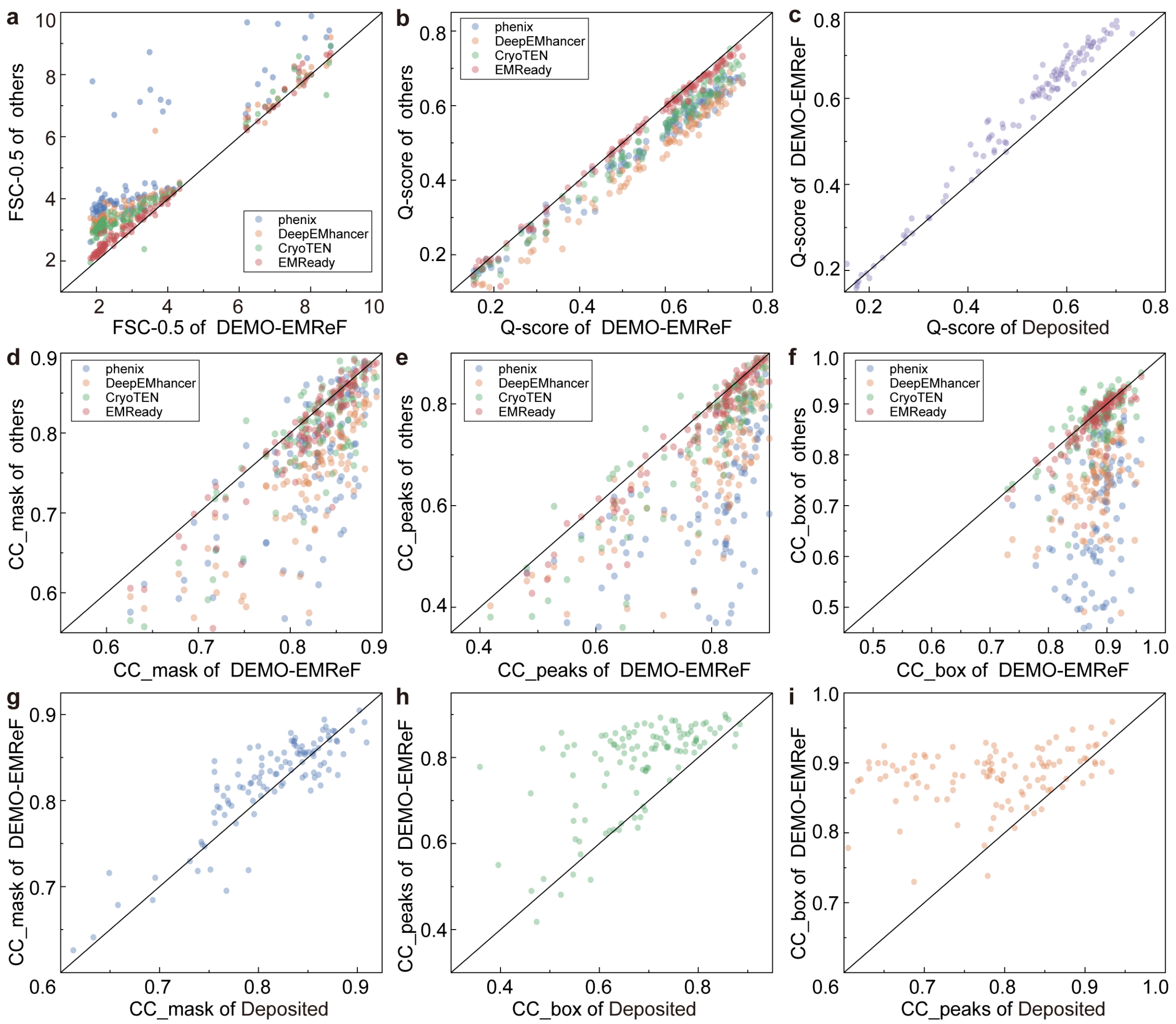


**Supplementary Figure 1 | Head-to-head comparison of DEMO-EMReF-enhanced maps with deposited maps and maps processed by other methods on the primary-map dataset.** (**a**) Comparison of FSC-0.5 resolution between DEMO-EMReF and other methods. (**b**) Comparison of Q-scores between DEMO-EMReF and other methods. (**c**) Q-score comparison between DEMO-EMReF-enhanced maps and the deposited maps. (**d-f**) Comparisons of CC_mask, CC_peaks, and CC_box, respectively, between DEMO-EMReF and other methods. (**g-i**) Comparisons of CC_mask, CC_peaks, and CC_box, respectively, between DEMO-EMReF-enhanced maps and the deposited maps.

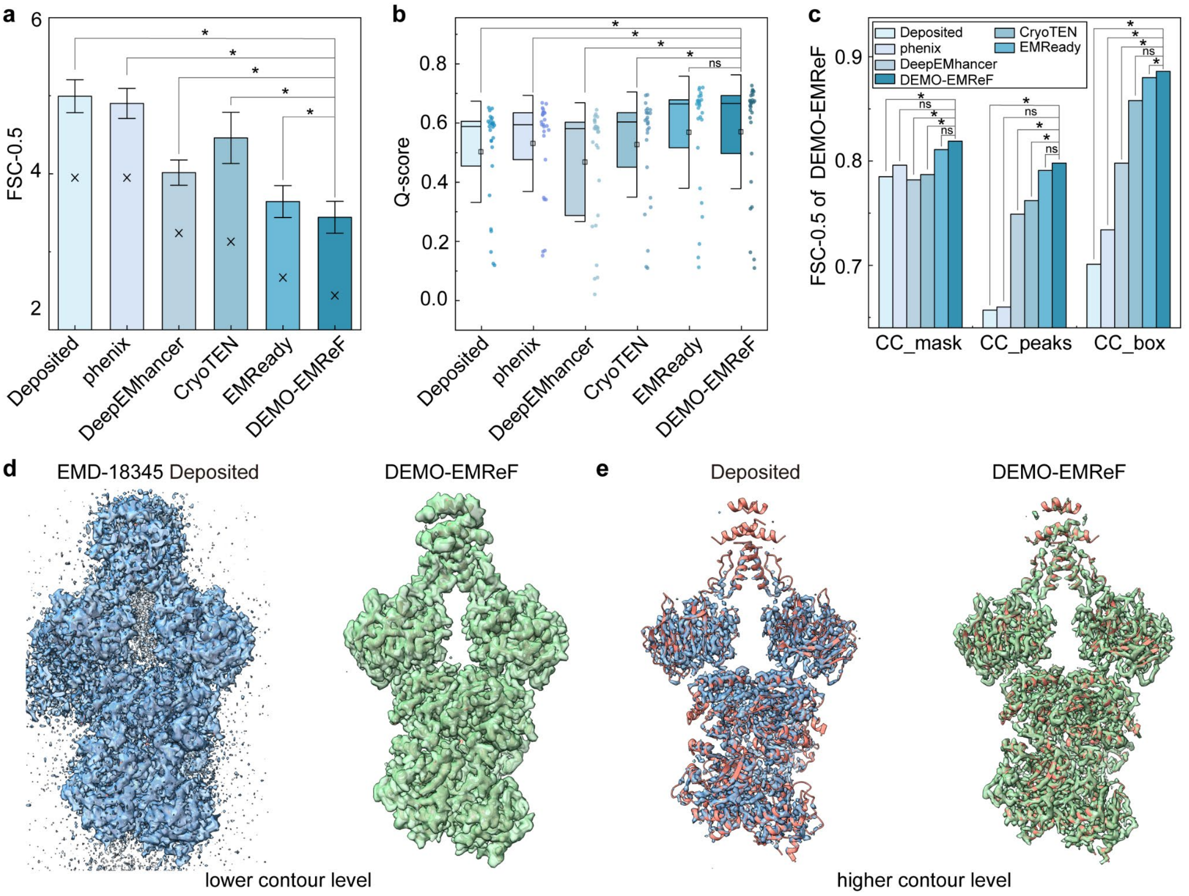


**Supplementary Figure 2 | Comparison of DEMO-EMReF-enhanced maps with deposited maps and maps processed by other methods on the half-map dataset. (a)** Bar chart of FSC-0.5 resolution for each refinement method, where bar heights represent the mean and crosses indicate the median. **(b)** Box plots of Q-scores showing local map quality for different methods, with squares representing the mean and horizontal lines indicating the median. **(c)** Bar plots of CC_mask, CC_box, and CC_peaks for different methods. **(d)** Comparison between the original averaged map of EMD-18345 and the corresponding DEMO-EMReF-enhanced map at a lower contour level. **(e)** Comparison between the original averaged map of EMD-18345 and the corresponding DEMO-EMReF-enhanced map at a higher contour level. Asterisks indicate statistically significant differences ($p$ < 0.0001, two-sided Wilcoxon signed-rank test compared with the baseline), whereas "ns" indicates no statistically significant difference.

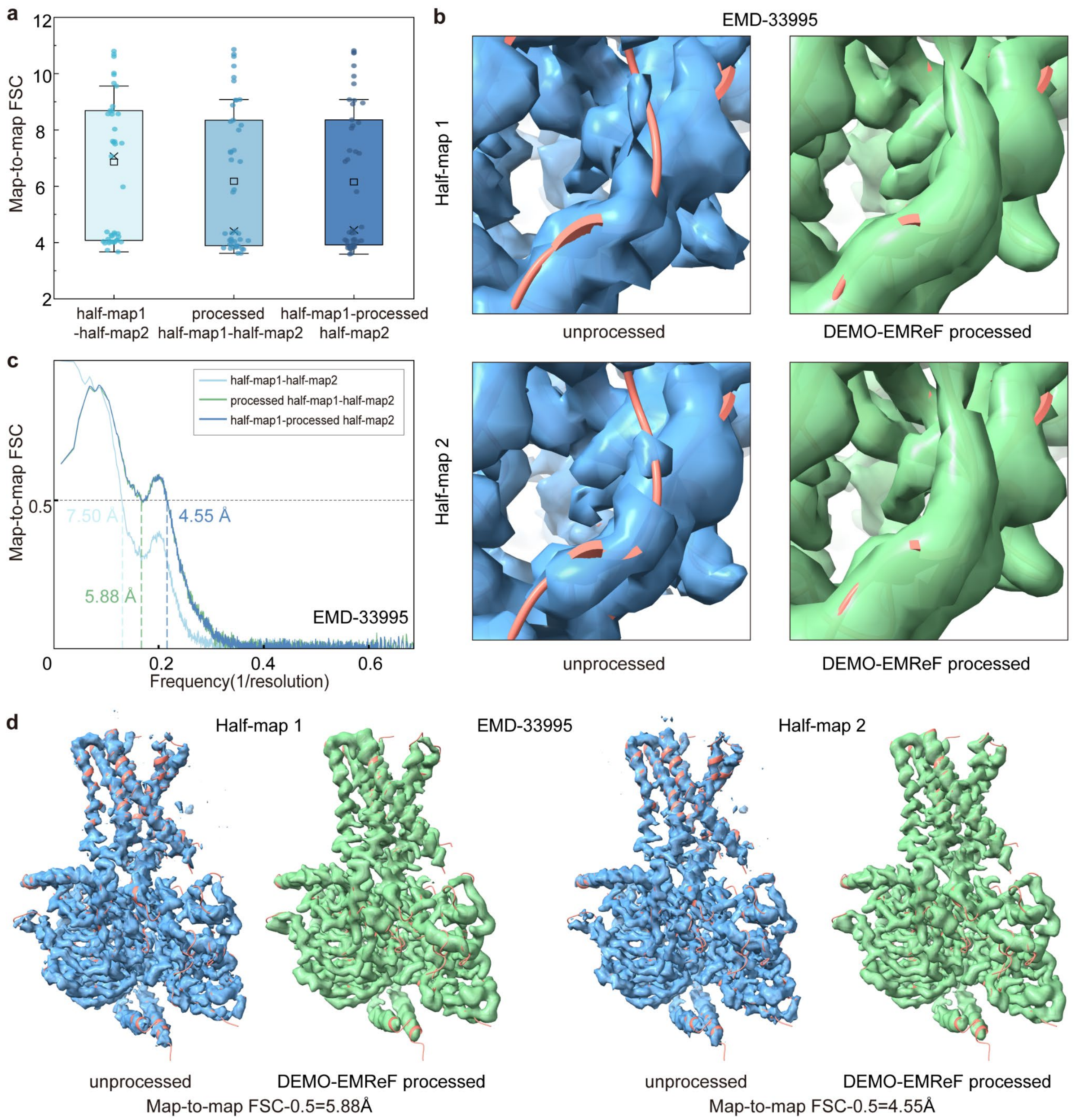


**Supplementary Figure 3 | Results of the cross-validation experiment on the half-map dataset.** **(a)** Box plots of unmasked FSC-0.5 resolution between the DEMO-EMReF-processed half-map and its corresponding unprocessed complementary half-map. Crosses and squares denote the median and mean, respectively, and box edges indicate the first and third quartiles. **(b)** Enlarged views of the unprocessed half-map and the DEMO-EMReF-processed half-map of EMD-33995 (PDB ID: 7YP7), with the top panel showing Half-map 1 and the bottom panel showing Half-map 2. **(c)** Unmasked FSC curves for unprocessed half-map pairs and for pairs consisting of one DEMO-EMReF-processed half-map and one unprocessed half-map (EMD-33995). **(d)** Global comparison between deposited half-maps and DEMO-EMReF-processed half-maps.

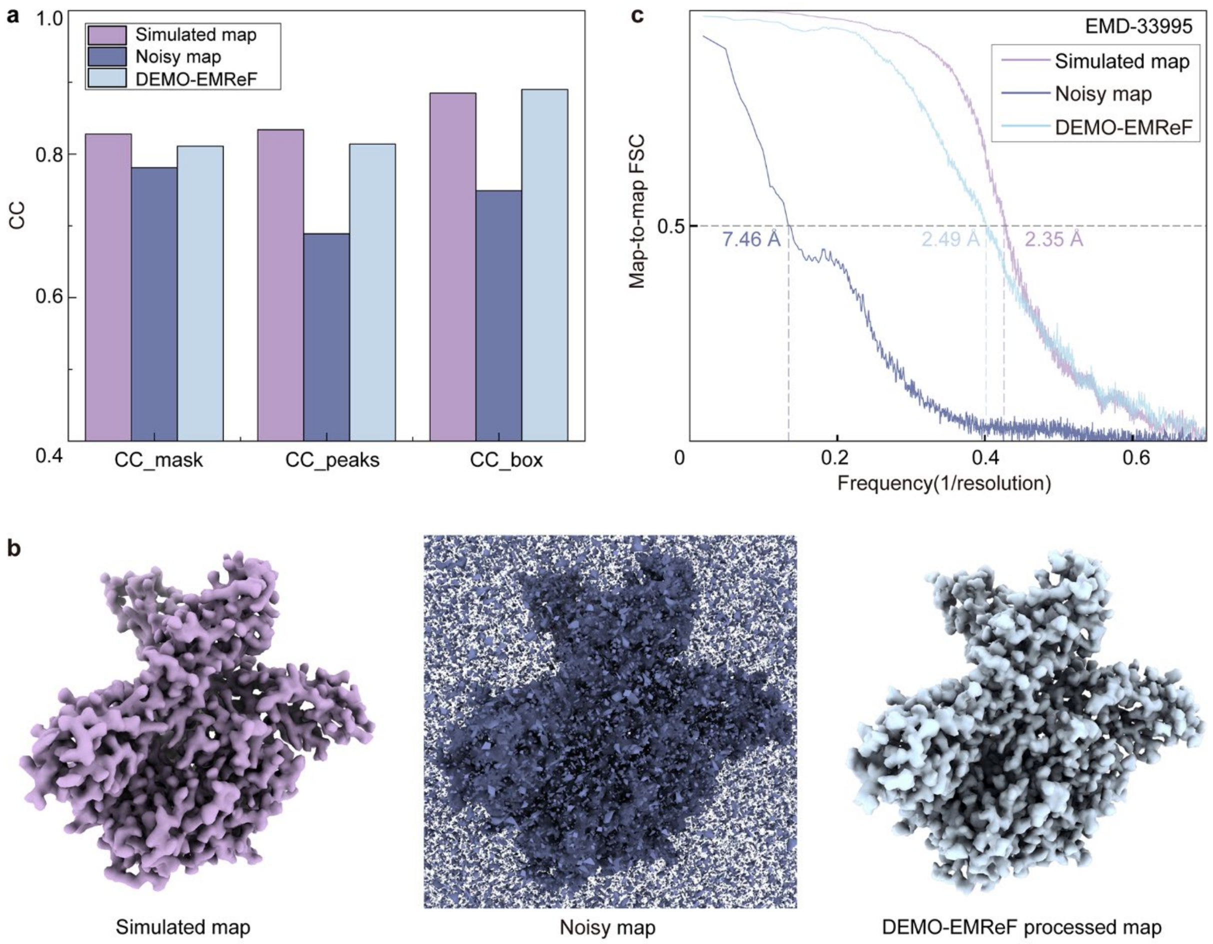


**Supplementary Figure 4 | Results showing DEMO-EMReF recovering the simulated map by identifying artificially added noise. (a)** Bar chart comparing the average CC values of the simulated maps, noisy maps, and DEMO-EMReF-processed maps on the primary-map dataset. **(b)** Visual comparison of the simulated map (left), the raw noisy map (center), and the DEMO-EMReF-processed map (right). The simulated map density was normalized to a maximum value of 1.0 prior to noise addition. Gaussian noise (mean = 0, $\sigma = 0.2$) was added to generate the noisy map. The density contour threshold of the DEMO-EMReF-processed map was adjusted to enclose a volume equivalent to that of the simulated and noisy maps for unbiased comparison. **(c)** Unmasked map-to-map FSC curves comparing the simulated map, the noisy map, and the noisy map after DEMO-EMReF processing.

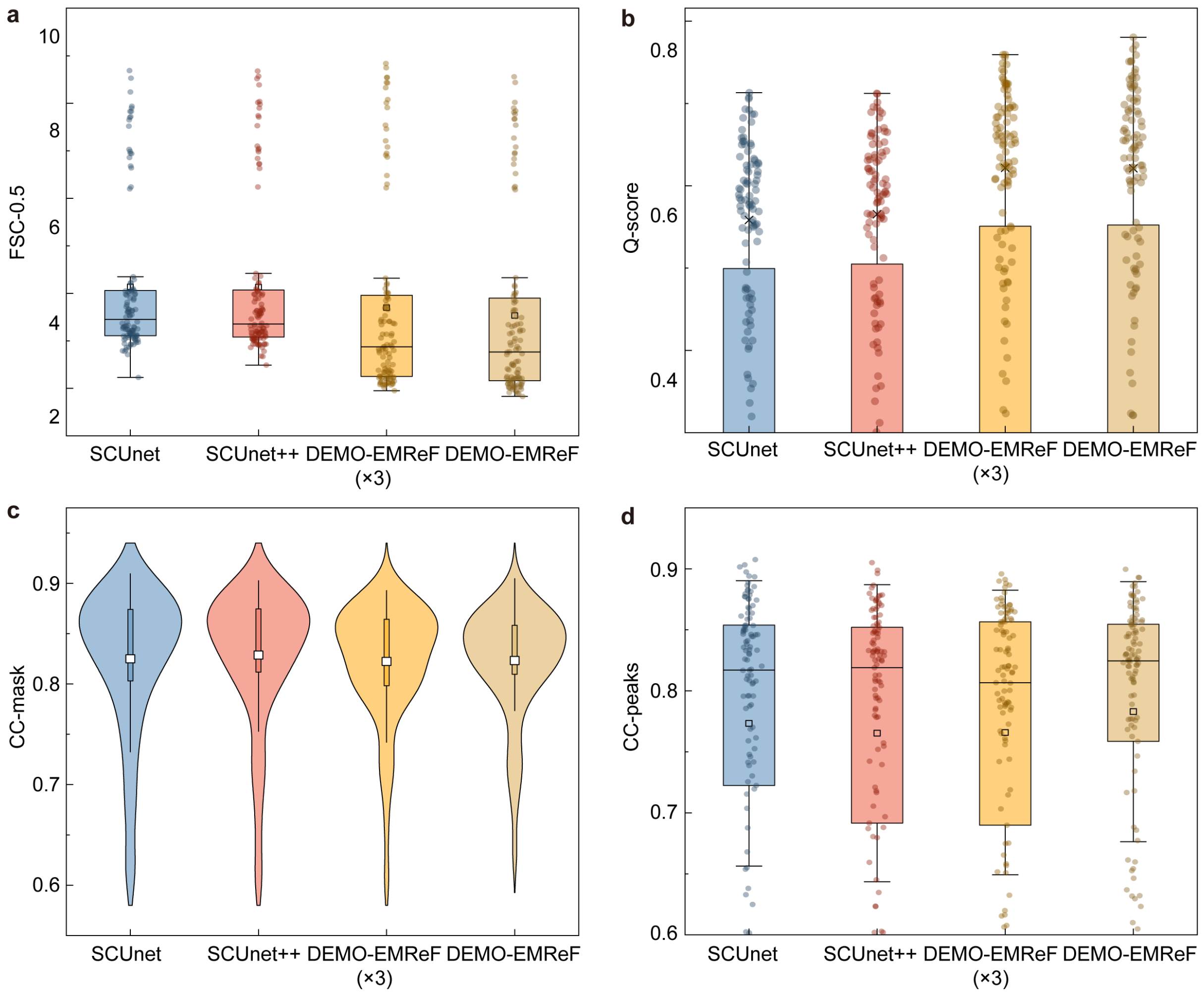


**Supplementary Figure 5 | Results of ablation experiments. (a)** Box plots showing the distribution of map-model FSC values (0.5 cutoff) for each ablated variant and the baseline. The median is indicated by the center line, the mean by a square symbol, and individual map results by circles. **(b)** Q-scores for each ablated variant and the baseline, shown as bar plots with overlaid scatter points representing individual cases. **(c,d)** Distributions of CC_mask (c) and CC_peaks (d), shown as violin plots (left) and box plots (right), with squares indicating the mean and horizontal lines indicating the median.

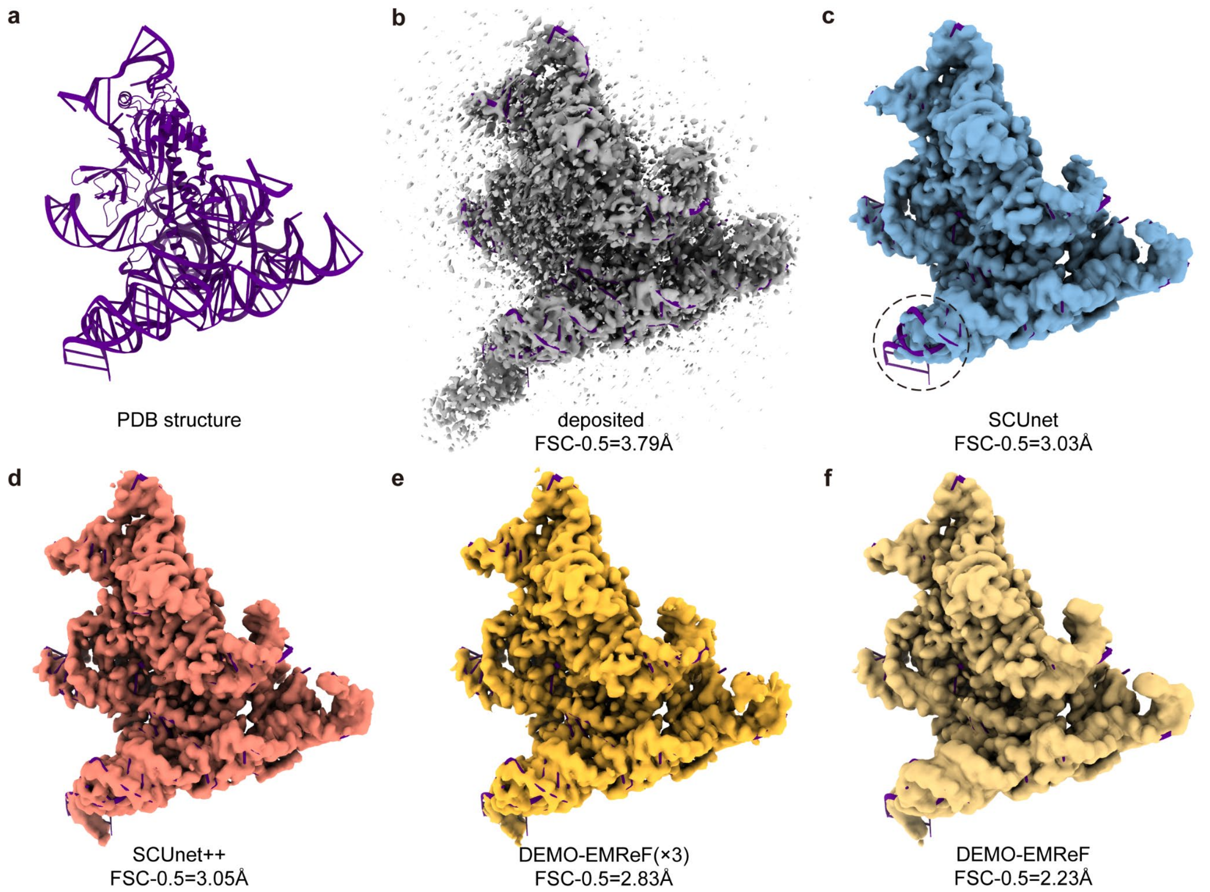


**Supplementary Figure 6 | Representative example comparing maps processed by different ablated variants.** **(a)** Deposited atomic structure (PDB: 8DMB). **(b)** Original deposited density map of EMD-27533. **(c-f)** Density maps produced by different ablated variants of DEMO-EMReF. The dashed circle highlights the nucleic acid density region for comparison.

### Supplementary Tables

**Supplementary Table 1 | Average results of different methods on the primary-map dataset.** Values were calculated only over test cases for which all methods produced valid results. Bold values indicate the best performance for each metric.

| Methods | FSC-0.5 (Å) | Q-score | CC_mask | CC_box | CC_peaks |
|---|---|---|---|---|---|
| Deposited | 5.03 | 0.498 | 0.810 | 0.785 | 0.681 |
| phenix | 5.03 | 0.486 | 0.757 | 0.676 | 0.586 |
| DeepEMhancer | 4.67 | 0.442 | 0.738 | 0.742 | 0.658 |
| CryoTEN | 4.45 | 0.501 | 0.794 | 0.868 | 0.735 |
| EMReady | 3.88 | 0.537 | 0.806 | 0.872 | 0.761 |
| **DEMO-EMReF** | **3.52** | **0.549** | **0.822** | **0.882** | **0.781** |

**Supplementary Table 2 | Summary of map-to-model FSC-0.5 resolution for each case in the primary-map dataset.** A dash ("-") indicates that the method failed to process the case or that phenix.mtriage did not return a valid FSC-0.5 value.

| FSC-0.5 | | | | | | | |
|---|---|---|---|---|---|---|---|
| EMDBID | PDBID | Deposited | phenix | DeepEMhancer | CryoTEN | EMReady | DEMO-EMReF |
| EMD-3605 | 5N9Y | 4.29 | 4.29 | 4.25 | 4.22 | 4.14 | 4.14 |
| EMD-8745 | 5VY9 | 6.98 | 6.98 | 7 | 6.97 | 6.94 | 6.96 |
| EMD-6714 | 5XB1 | 3.04 | 3.04 | 3.03 | 2.98 | 2.12 | 2.02 |
| EMD-7065 | 6B7Y | 10.01 | 9.88 | 8.24 | 7.99 | 8.01 | 8.02 |
| EMD-3896 | 6EMK | 9.4 | 9.42 | 8.46 | 8.37 | 8.61 | 8.52 |
| EMD-0088 | 6GY6 | 4.07 | 4.07 | - | 3.89 | 3.33 | 3.32 |
| EMD-9915 | 6K4M | 7.91 | 7.91 | - | 7.45 | 6.95 | 6.94 |
| EMD-20551 | 6Q0K | 7.94 | 8.09 | 7.63 | 7.89 | 7.8 | 7.67 |
| EMD-4975 | 6ROW | 4.3 | 4.31 | 4.23 | 4.22 | 4.19 | 4.17 |
| EMD-20724 | 6UC0 | 15.77 | 15.79 | 16.09 | 7.71 | 8.36 | 7.77 |
| EMD-21913 | 6WUH | 3.62 | 3.71 | 3.63 | 3.53 | 3.13 | 3.05 |
| EMD-11439 | 6ZU9 | 9.95 | 9.96 | 8.05 | 8.59 | 8.21 | 7.67 |
| EMD-12054 | 7B6H | 9.56 | 9.2 | 8.28 | 7.34 | 8.64 | 8.44 |
| EMD-30342 | 7CEC | 4.54 | 4.31 | 4.17 | 4.02 | 4 | 3.97 |
| EMD-30346 | 7CFS | 3.64 | 3.67 | 3.76 | 3.46 | 2.93 | 2.99 |
| EMD-30384 | 7CKB | 3.26 | 3.26 | - | 3.16 | 2.24 | 2.07 |
| EMD-30535 | 7D0I | 3.2 | 3.2 | 3.23 | 2.92 | 2.21 | 2.04 |
| EMD-31305 | 7EU0 | 3.37 | 3.37 | 3.33 | 3.15 | 2.43 | 2.21 |
| EMD-31561 | 7FER | 4.35 | 4.35 | 4.05 | 4.04 | 3.83 | 4 |
| EMD-31595 | 7FIF | 6.94 | 7.01 | 6.88 | 6.54 | 6.62 | 6.25 |
| EMD-22359 | 7JK2 | 3.71 | 3.71 | - | 3.08 | 2.48 | 2.21 |
| EMD-22375 | 7JLO | 3.82 | 3.82 | 3.29 | 2.98 | 2.2 | 1.95 |
| EMD-22445 | 7JRG | 4.29 | 4.29 | 3.16 | 3.09 | 2.34 | 2.06 |
| EMD-22233 | 7KEU | 6.01 | 4.37 | 6.19 | 4.05 | 3.47 | 3.65 |
| EMD-22879 | 7KHF | 4.41 | 4.41 | 3.7 | 3.52 | 2.83 | 2.81 |
| EMD-22901 | 7KJX | 3.82 | 3.83 | 3.28 | 3.18 | 2.7 | 2.49 |
| EMD-23340 | 7LHH | 8.91 | 8.91 | 9.2 | 8.93 | 8.7 | 8.56 |
| EMD-23526 | 7LUP | 7.97 | 9.68 | 6.31 | 6.29 | 6.21 | 6.22 |
| EMD-23930 | 7MP6 | 9.18 | 9.18 | 7.78 | - | 7.84 | 7.84 |
| EMD-24073 | 7MXL | 3.92 | 3.93 | - | 3.2 | 2.83 | 2.56 |
| EMD-12296 | 7NF6 | 3.19 | 3.19 | 3.33 | 2.8 | 2.48 | 2.48 |
| EMD-12309 | 7NG8 | 3.36 | 3.36 | 3.33 | 3.17 | 2.23 | 1.93 |
| EMD-12600 | 7NUR | 3.2 | 3.22 | 3.28 | 3.03 | 2.29 | 2.02 |
| EMD-12808 | 7OCI | 3.41 | 3.42 | 3.43 | 3.19 | 2.45 | 2.21 |
| EMD-12993 | 7ON1 | 3.49 | 3.49 | 3.37 | 3.16 | 2.43 | 2.18 |
| EMD-13035 | 7OQZ | 12.88 | 12.69 | 3.83 | 3.58 | 2.71 | 2.33 |
| EMD-13358 | 7PEV | 6.61 | 6.73 | 6.58 | 6.41 | 6.29 | 6.18 |

| | | | | | | | |
|---|---|---|---|---|---|---|---|
| EMD-13587 | 7PQ5 | 3.58 | 3.6 | 3.23 | 2.94 | 2.21 | 1.98 |
| EMD-13827 | 7Q56 | 7.15 | 7.15 | 6.99 | 6.86 | 6.83 | 6.83 |
| EMD-14145 | 7QTQ | 3.67 | 3.67 | - | 3.35 | 3.13 | 2.88 |
| EMD-14244 | 7R21 | 3.93 | 3.81 | 3.28 | 3.2 | 2.65 | 2.52 |
| EMD-24473 | 7RI4 | 4.12 | 4.13 | 3.84 | 3.62 | 3.41 | 3.35 |
| EMD-24889 | 7S82 | 4.01 | 4.01 | - | 3.62 | 3.41 | 3.17 |
| EMD-24991 | 7SBX | 3.68 | 3.67 | 2.95 | 2.64 | 2.08 | 1.89 |
| EMD-26377 | 7U6O | 3.87 | 3.87 | 3.8 | 3.66 | 3.42 | 3.3 |
| EMD-26512 | 7UHC | 4.14 | 4.14 | 3.4 | 3.19 | 2.55 | 2.24 |
| EMD-32536 | 7WIU | 6.7 | 6.7 | 3.71 | 3.55 | 2.69 | 2.5 |
| EMD-32845 | 7WV4 | 7.2 | 7.19 | - | 4 | 3.84 | 3.8 |
| EMD-32894 | 7WYU | 3.92 | 3.92 | 3.39 | 3.19 | 2.56 | 2.32 |
| EMD-33180 | 7XFZ | 3.08 | 3.08 | 3.14 | 2.87 | 2.41 | 2.24 |
| EMD-33192 | 7XHA | 3.83 | 3.84 | 3.5 | 3.39 | 2.8 | 2.57 |
| EMD-33666 | 7Y7I | 3.68 | 3.68 | 3.29 | 3.3 | 2.56 | 2.15 |
| EMD-33995 | 7YP7 | 3.27 | 3.28 | 3.25 | 3.08 | 2.68 | 2.4 |
| EMD-14849 | 7ZOX | 4.15 | 4.19 | 4.26 | 3.96 | 3.84 | 3.83 |
| EMD-15330 | 8AC1 | 6.84 | 6.81 | 4.14 | 4.08 | 4 | 3.86 |
| EMD-16555 | 8CC6 | 3.56 | 3.56 | 3.25 | 3 | 2.26 | 2.18 |
| EMD-26978 | 8CT2 | 7.85 | 7.78 | 3.02 | 2.9 | 2.1 | 1.89 |
| EMD-27034 | 8CX2 | 4.04 | 3.94 | 3.49 | 3.25 | 2.95 | 2.91 |
| EMD-27104 | 8D0B | 4.27 | 4.27 | 3.81 | 3.62 | 3.14 | 2.84 |
| EMD-27178 | 8D49 | 3.79 | 3.78 | 3.35 | 3.23 | 2.39 | 2.11 |
| EMD-27533 | 8DMB | 3.79 | 3.73 | 3.05 | 2.98 | 2.95 | 2.23 |
| EMD-28161 | 8EIQ | 3.75 | 3.67 | 2.89 | 2.76 | 2.15 | 2 |
| EMD-28204 | 8EKI | 7.69 | 7.69 | 7.03 | 6.73 | 6.5 | 6.52 |
| EMD-28534 | 8EPX | 3.58 | 3.58 | 3.13 | 3.06 | 2.49 | 2.09 |
| EMD-29686 | 8G30 | 3.59 | 3.52 | 3.33 | 3.25 | 3.18 | 2.96 |
| EMD-29936 | 8GCN | 4.43 | 4.48 | 4.53 | 4.48 | 4.32 | 4.33 |
| EMD-35304 | 8IAJ | 3.61 | 3.61 | 3.1 | 2.98 | 2.25 | 2.03 |
| EMD-35728 | 8IUO | 7.24 | 7.11 | 4.32 | 4.15 | 4.03 | 4.02 |
| EMD-35912 | 8J12 | 3.07 | 3.07 | 3.15 | 3.17 | 2.87 | 2.1 |
| EMD-35948 | 8J2F | 3.15 | 3.15 | 3.1 | 3.02 | 2.3 | 2.2 |
| EMD-36046 | 8J7R | 4.13 | 4.13 | 3.44 | 3.48 | 3.36 | 3.15 |
| EMD-36070 | 8J8H | 4.12 | 4.12 | 3.92 | 3.86 | 3.76 | 3.65 |
| EMD-36824 | 8K21 | 4.06 | 4.05 | 4.02 | 3.8 | 3.3 | 3.2 |
| EMD-17341 | 8P0X | 9.97 | 10.03 | 8.08 | - | 7.65 | 7.81 |
| EMD-17632 | 8PEN | 3.5 | 3.5 | 3.95 | 3.28 | 3.05 | 2.77 |
| EMD-17979 | 8PVX | 3.72 | 3.72 | 15.69 | 3.7 | 3.44 | 3.23 |
| EMD-18345 | 8QE8 | 3.85 | 3.85 | 3.9 | 3.36 | 2.53 | 2.22 |
| EMD-18453 | 8QJX | 3.52 | 3.53 | 3.05 | 2.96 | 2.18 | 1.91 |
| EMD-18471 | 8QKU | 4 | 4 | 3.86 | 3.62 | 3.06 | 3.01 |
| EMD-18608 | 8QQX | 3.05 | 3.05 | 3.1 | 2.74 | 2.17 | 2.01 |

| | | | | | | | |
|---|---|---|---|---|---|---|---|
| EMD-18638 | 8QSP | 4.54 | 4.5 | 4.05 | 3.97 | 3.61 | 3.36 |
| EMD-18778 | 8QZM | 3.49 | 3.49 | 3.18 | 3.08 | 2.23 | 2.03 |
| EMD-40241 | 8S9P | 4.09 | 4.15 | 4.08 | 3.98 | 4 | 3.94 |
| EMD-41011 | 8T3T | 3.59 | 3.58 | 3.11 | 2.88 | 2.23 | 2.03 |
| EMD-41355 | 8TKO | 3.94 | 3.95 | 3.28 | 3.08 | 2.82 | 2.39 |
| EMD-42774 | 8UXQ | 8.24 | 8.24 | 8.27 | 8.17 | 7.4 | 7.54 |
| EMD-42795 | 8UY8 | 3.15 | 3.15 | 3.14 | 2.94 | 2.21 | 2.08 |
| EMD-42960 | 8V3Y | 3.73 | 3.73 | - | 2.38 | 3.55 | 3.34 |
| EMD-43196 | 8VG0 | 3.72 | 3.73 | 3.2 | 3.02 | 2.29 | 1.98 |
| EMD-43304 | 8VK4 | 3.4 | 3.4 | 3.45 | 3.22 | 2.36 | 2.1 |
| EMD-43316 | 8VKH | 8.7 | 8.72 | 4.02 | 3.82 | 3.53 | 3.49 |
| EMD-43346 | 8VLW | 4 | 4.02 | 3.42 | 3.32 | 2.34 | 2.17 |
| EMD-37439 | 8WCE | 3.8 | 3.8 | 3.21 | 3.16 | 2.33 | 2.09 |
| EMD-37658 | 8WMQ | 3.87 | 3.88 | 3.29 | 3.21 | 2.54 | 2.09 |
| EMD-38025 | 8X31 | 7.5 | 7.5 | 7.53 | 7.5 | 7.54 | 7.27 |
| EMD-39454 | 8YON | 9.68 | 9.63 | 7.23 | 7.21 | 7.04 | 7.08 |
| EMD-60471 | 8ZTS | 3.67 | 3.68 | 3.51 | 3.28 | 2.55 | 2.21 |
| EMD-44441 | 9BCX | 8.53 | 8.53 | 7.98 | 7.95 | 7.91 | 7.9 |
| EMD-45805 | 9CPO | 3.92 | 3.92 | 3.59 | 3.46 | 2.9 | 2.41 |
| EMD-45924 | 9CU0 | 7.51 | 7.51 | 4.1 | 4.02 | 3.7 | 3.51 |
| EMD-45988 | 9CXF | 3.35 | 3.34 | 3.13 | 3.01 | 3.08 | 2.73 |
| EMD-46645 | 9D8P | 4.09 | 4.08 | 3.33 | 3.31 | 2.84 | 2.76 |
| EMD-47091 | 9DOU | 4.01 | 4 | 3.04 | 2.9 | 2.33 | 2.14 |
| EMD-47476 | 9E3A | 3.64 | 3.62 | 20.57 | 37.49 | 21.05 | 3.37 |
| EMD-50262 | 9F9X | 2.22 | 2.5 | - | 2.66 | 2.52 | 2.25 |
| EMD-50289 | 9FB6 | 1.78 | 2.61 | 3.38 | 1.95 | 2.09 | 1.83 |
| EMD-50522 | 9FKD | 3.73 | 3.73 | 3.51 | 3.43 | 2.87 | 2.71 |
| EMD-51241 | 9GD1 | 4.49 | 4.5 | 4.16 | 3.98 | 3.61 | 3.61 |
| EMD-51918 | 9H82 | - | 18.74 | 12.79 | 21.56 | 12.88 | 12.49 |
| EMD-61073 | 9J1J | 7.12 | 7.12 | 3.85 | 3.61 | 3.34 | 3.22 |
| EMD-62812 | 9L4F | 8.03 | 8.14 | 6.44 | 6.97 | 6.95 | 6.72 |

**Supplementary Table 3 | Summary of Q-score for each case in the primary-map dataset.** A dash ("-") indicates that the method failed to process the case.

| Q-score | | | | | | | |
|---|---|---|---|---|---|---|---|
| EMDBID | PDBID | Deposited | phenix | DeepEMhancer | CryoTEN | EMReady | DEMO-EMReF |
| EMD-3605 | 5N9Y | 0.416 | 0.428 | 0.364 | 0.401 | 0.470 | 0.470 |
| EMD-8745 | 5VY9 | 0.201 | 0.188 | 0.059 | 0.183 | 0.189 | 0.182 |
| EMD-6714 | 5XB1 | 0.701 | 0.662 | 0.659 | 0.681 | 0.732 | 0.780 |
| EMD-7065 | 6B7Y | 0.195 | 0.166 | 0.078 | 0.175 | 0.188 | 0.194 |
| EMD-3896 | 6EMK | 0.179 | 0.161 | 0.121 | 0.165 | 0.180 | 0.175 |
| EMD-0088 | 6GY6 | 0.439 | 0.463 | - | 0.457 | 0.546 | 0.545 |
| EMD-9915 | 6K4M | 0.286 | 0.334 | - | 0.258 | 0.317 | 0.321 |
| EMD-20551 | 6Q0K | 0.274 | 0.233 | 0.184 | 0.231 | 0.276 | 0.278 |
| EMD-4975 | 6ROW | 0.463 | 0.456 | 0.426 | 0.463 | 0.495 | 0.500 |
| EMD-20724 | 6UC0 | 0.184 | 0.173 | 0.112 | 0.178 | 0.181 | 0.189 |
| EMD-21913 | 6WUH | 0.582 | 0.577 | 0.537 | 0.586 | 0.622 | 0.622 |
| EMD-11439 | 6ZU9 | 0.270 | 0.265 | 0.238 | 0.239 | 0.268 | 0.282 |
| EMD-12054 | 7B6H | 0.322 | 0.259 | 0.241 | 0.282 | 0.290 | 0.321 |
| EMD-30342 | 7CEC | 0.422 | 0.428 | 0.338 | 0.416 | 0.437 | 0.430 |
| EMD-30346 | 7CFS | 0.583 | 0.579 | 0.526 | 0.597 | 0.637 | 0.646 |
| EMD-30384 | 7CKB | 0.702 | 0.675 | - | 0.717 | 0.753 | 0.766 |
| EMD-30535 | 7D0I | 0.628 | 0.626 | 0.596 | 0.629 | 0.679 | 0.704 |
| EMD-31305 | 7EU0 | 0.580 | 0.569 | 0.543 | 0.629 | 0.615 | 0.622 |
| EMD-31561 | 7FER | 0.536 | 0.542 | 0.498 | 0.543 | 0.603 | 0.606 |
| EMD-31595 | 7FIF | 0.350 | 0.327 | 0.215 | 0.356 | 0.349 | 0.360 |
| EMD-22359 | 7JK2 | 0.578 | 0.603 | - | 0.600 | 0.644 | 0.644 |
| EMD-22375 | 7JLO | 0.643 | 0.629 | 0.598 | 0.655 | 0.707 | 0.731 |
| EMD-22445 | 7JRG | 0.650 | 0.642 | 0.621 | 0.669 | 0.706 | 0.721 |
| EMD-22233 | 7KEU | 0.368 | 0.317 | 0.325 | 0.348 | 0.423 | 0.436 |
| EMD-22879 | 7KHF | 0.598 | 0.577 | 0.563 | 0.609 | 0.650 | 0.658 |
| EMD-22901 | 7KJX | 0.615 | 0.612 | 0.568 | 0.631 | 0.671 | 0.680 |
| EMD-23340 | 7LHH | 0.209 | 0.187 | -0.089 | 0.184 | 0.189 | 0.210 |
| EMD-23526 | 7LUP | 0.271 | 0.267 | 0.185 | 0.258 | 0.274 | 0.264 |
| EMD-23930 | 7MP6 | 0.175 | 0.160 | 0.119 | 0.161 | 0.162 | 0.172 |
| EMD-24073 | 7MXL | 0.601 | 0.598 | - | 0.611 | 0.646 | 0.648 |
| EMD-12296 | 7NF6 | 0.664 | 0.651 | 0.631 | 0.677 | 0.702 | 0.718 |
| EMD-12309 | 7NG8 | 0.599 | 0.615 | 0.580 | 0.632 | 0.703 | 0.723 |
| EMD-12600 | 7NUR | 0.667 | 0.645 | 0.622 | 0.688 | 0.719 | 0.733 |
| EMD-12808 | 7OCI | 0.618 | 0.599 | 0.575 | 0.630 | 0.682 | 0.694 |
| EMD-12993 | 7ON1 | 0.564 | 0.547 | 0.501 | 0.578 | 0.609 | 0.633 |
| EMD-13035 | 7OQZ | 0.509 | 0.512 | 0.459 | 0.518 | 0.586 | 0.594 |
| EMD-13358 | 7PEV | 0.277 | 0.264 | 0.212 | 0.248 | 0.268 | 0.292 |
| EMD-13587 | 7PQ5 | 0.676 | 0.680 | 0.649 | 0.712 | 0.727 | 0.737 |

| | | | | | | | |
|---|---|---|---|---|---|---|---|
| EMD-13827 | 7Q56 | 0.288 | 0.279 | 0.207 | 0.281 | 0.270 | 0.286 |
| EMD-14145 | 7QTQ | 0.603 | 0.593 | - | 0.606 | 0.648 | 0.643 |
| EMD-14244 | 7R21 | 0.564 | 0.551 | 0.500 | 0.553 | 0.591 | 0.601 |
| EMD-24473 | 7RI4 | 0.532 | 0.504 | 0.486 | 0.534 | 0.564 | 0.555 |
| EMD-24889 | 7S82 | 0.557 | 0.539 | - | 0.532 | 0.612 | 0.605 |
| EMD-24991 | 7SBX | 0.682 | 0.671 | 0.664 | 0.708 | 0.733 | 0.752 |
| EMD-26377 | 7U6O | 0.449 | 0.466 | 0.404 | 0.441 | 0.514 | 0.515 |
| EMD-26512 | 7UHC | 0.560 | 0.534 | 0.529 | 0.586 | 0.620 | 0.634 |
| EMD-32536 | 7WIU | 0.575 | 0.566 | 0.538 | 0.583 | 0.653 | 0.662 |
| EMD-32845 | 7WV4 | 0.431 | 0.437 | - | 0.414 | 0.455 | 0.466 |
| EMD-32894 | 7WYU | 0.545 | 0.530 | 0.511 | 0.561 | 0.604 | 0.607 |
| EMD-33180 | 7XFZ | 0.651 | 0.639 | 0.602 | 0.666 | 0.682 | 0.690 |
| EMD-33192 | 7XHA | 0.552 | 0.548 | 0.509 | 0.562 | 0.619 | 0.618 |
| EMD-33666 | 7Y7I | 0.543 | 0.549 | 0.503 | 0.563 | 0.585 | 0.610 |
| EMD-33995 | 7YP7 | 0.636 | 0.612 | 0.596 | 0.653 | 0.683 | 0.690 |
| EMD-14849 | 7ZOX | 0.478 | 0.439 | 0.383 | 0.456 | 0.502 | 0.496 |
| EMD-15330 | 8AC1 | 0.408 | 0.314 | 0.353 | 0.389 | 0.418 | 0.410 |
| EMD-16555 | 8CC6 | 0.644 | 0.636 | 0.605 | 0.670 | 0.712 | 0.741 |
| EMD-26978 | 8CT2 | 0.690 | 0.631 | 0.637 | 0.660 | 0.737 | 0.751 |
| EMD-27034 | 8CX2 | 0.453 | 0.440 | 0.403 | 0.446 | 0.468 | 0.475 |
| EMD-27104 | 8D0B | 0.503 | 0.463 | 0.451 | 0.504 | 0.544 | 0.536 |
| EMD-27178 | 8D49 | 0.601 | 0.620 | 0.576 | 0.629 | 0.680 | 0.692 |
| EMD-27533 | 8DMB | 0.624 | 0.607 | 0.593 | 0.639 | 0.641 | 0.687 |
| EMD-28161 | 8EIQ | 0.692 | 0.675 | 0.668 | 0.710 | 0.757 | 0.771 |
| EMD-28204 | 8EKI | 0.227 | 0.189 | 0.129 | 0.218 | 0.230 | 0.229 |
| EMD-28534 | 8EPX | 0.649 | 0.633 | 0.625 | 0.667 | 0.695 | 0.720 |
| EMD-29686 | 8G30 | 0.588 | 0.635 | 0.570 | 0.622 | 0.659 | 0.658 |
| EMD-29936 | 8GCN | 0.357 | 0.345 | 0.289 | 0.352 | 0.390 | 0.398 |
| EMD-35304 | 8IAJ | 0.676 | 0.665 | 0.645 | 0.700 | 0.736 | 0.759 |
| EMD-35728 | 8IUO | 0.444 | 0.441 | 0.392 | 0.439 | 0.503 | 0.510 |
| EMD-35912 | 8J12 | 0.570 | 0.573 | 0.021 | 0.578 | 0.611 | 0.653 |
| EMD-35948 | 8J2F | 0.643 | 0.629 | 0.611 | 0.671 | 0.709 | 0.723 |
| EMD-36046 | 8J7R | 0.492 | 0.477 | 0.436 | 0.503 | 0.528 | 0.538 |
| EMD-36070 | 8J8H | 0.475 | 0.476 | 0.425 | 0.473 | 0.498 | 0.498 |
| EMD-36824 | 8K21 | 0.442 | 0.475 | 0.433 | 0.472 | 0.552 | 0.549 |
| EMD-17341 | 8P0X | 0.320 | 0.310 | 0.261 | 0.286 | 0.326 | 0.323 |
| EMD-17632 | 8PEN | 0.544 | 0.523 | 0.504 | 0.561 | 0.613 | 0.621 |
| EMD-17979 | 8PVX | 0.538 | 0.562 | 0.522 | 0.556 | 0.632 | 0.649 |
| EMD-18345 | 8QE8 | 0.633 | 0.587 | 0.604 | 0.626 | 0.691 | 0.716 |
| EMD-18453 | 8QJX | 0.589 | 0.592 | 0.577 | 0.593 | 0.651 | 0.664 |
| EMD-18471 | 8QKU | 0.442 | 0.444 | 0.383 | 0.460 | 0.505 | 0.505 |
| EMD-18608 | 8QQX | 0.655 | 0.671 | 0.635 | 0.693 | 0.723 | 0.731 |
| EMD-18638 | 8QSP | 0.471 | 0.459 | 0.430 | 0.477 | 0.533 | 0.532 |

| | | | | | | | |
|---|---|---|---|---|---|---|---|
| EMD-18778 | 8QZM | 0.608 | 0.633 | 0.602 | 0.637 | 0.664 | 0.706 |
| EMD-40241 | 8S9P | 0.477 | 0.472 | 0.417 | 0.459 | 0.483 | 0.476 |
| EMD-41011 | 8T3T | 0.641 | 0.639 | 0.613 | 0.667 | 0.685 | 0.702 |
| EMD-41355 | 8TKO | 0.594 | 0.609 | 0.570 | 0.627 | 0.672 | 0.683 |
| EMD-42774 | 8UXQ | 0.176 | 0.149 | 0.106 | 0.157 | 0.183 | 0.165 |
| EMD-42795 | 8UY8 | 0.654 | 0.628 | 0.612 | 0.676 | 0.718 | 0.729 |
| EMD-42960 | 8V3Y | 0.551 | 0.554 | - | 0.571 | 0.615 | 0.625 |
| EMD-43196 | 8VG0 | 0.579 | 0.583 | 0.545 | 0.597 | 0.611 | 0.655 |
| EMD-43304 | 8VK4 | 0.531 | 0.521 | 0.469 | 0.547 | 0.602 | 0.604 |
| EMD-43316 | 8VKH | 0.470 | 0.479 | 0.429 | 0.474 | 0.534 | 0.541 |
| EMD-43346 | 8VLW | 0.552 | 0.549 | 0.521 | 0.576 | 0.671 | 0.678 |
| EMD-37439 | 8WCE | 0.611 | 0.601 | 0.552 | 0.620 | 0.654 | 0.669 |
| EMD-37658 | 8WMQ | 0.591 | 0.578 | 0.550 | 0.600 | 0.657 | 0.672 |
| EMD-38025 | 8X31 | 0.155 | 0.158 | 0.066 | 0.146 | 0.115 | 0.216 |
| EMD-39454 | 8YON | 0.166 | 0.156 | 0.081 | 0.132 | 0.120 | 0.153 |
| EMD-60471 | 8ZTS | 0.603 | 0.574 | 0.586 | 0.610 | 0.687 | 0.699 |
| EMD-44441 | 9BCX | 0.165 | 0.141 | 0.078 | 0.143 | 0.144 | 0.152 |
| EMD-45805 | 9CPO | 0.573 | 0.565 | 0.531 | 0.583 | 0.611 | 0.625 |
| EMD-45924 | 9CU0 | 0.453 | 0.443 | 0.398 | 0.435 | 0.479 | 0.480 |
| EMD-45988 | 9CXF | 0.615 | 0.619 | 0.564 | 0.637 | 0.627 | 0.634 |
| EMD-46645 | 9D8P | 0.626 | 0.635 | 0.609 | 0.629 | 0.684 | 0.707 |
| EMD-47091 | 9DOU | 0.635 | 0.607 | 0.563 | 0.644 | 0.649 | 0.659 |
| EMD-47476 | 9E3A | 0.579 | 0.588 | 0.551 | 0.570 | 0.643 | 0.667 |
| EMD-50262 | 9F9X | 0.570 | 0.559 | - | 0.582 | 0.594 | 0.619 |
| EMD-50289 | 9FB6 | 0.733 | 0.671 | 0.690 | 0.752 | 0.732 | 0.751 |
| EMD-50522 | 9FKD | 0.588 | 0.569 | 0.546 | 0.583 | 0.653 | 0.655 |
| EMD-51241 | 9GD1 | 0.405 | 0.425 | 0.370 | 0.417 | 0.483 | 0.493 |
| EMD-51918 | 9H82 | 0.174 | 0.157 | 0.124 | 0.140 | 0.166 | 0.159 |
| EMD-61073 | 9J1J | 0.534 | 0.555 | 0.503 | 0.542 | 0.605 | 0.611 |
| EMD-62812 | 9L4F | 0.355 | 0.320 | 0.295 | 0.343 | 0.380 | 0.373 |

**Supplementary Table 4 | Summary of CC_mask for each case in the primary-map dataset.** A dash ("-") indicates that the method failed to process the case.

| CC_mask | | | | | | | |
|---|---|---|---|---|---|---|---|
| EMDBID | PDBID | Deposited | phenix | DeepEMhancer | CryoTEN | EMReady | DEMO-EMReF |
| EMD-3605 | 5N9Y | 0.801 | 0.797 | 0.680 | 0.835 | 0.826 | 0.827 |
| EMD-8745 | 5VY9 | 0.731 | 0.695 | 0.623 | 0.720 | 0.717 | 0.730 |
| EMD-6714 | 5XB1 | 0.879 | 0.880 | 0.866 | 0.893 | 0.891 | 0.870 |
| EMD-7065 | 6B7Y | 0.842 | 0.678 | 0.508 | 0.782 | 0.835 | 0.858 |
| EMD-3896 | 6EMK | 0.790 | 0.654 | 0.634 | 0.653 | 0.733 | 0.719 |
| EMD-0088 | 6GY6 | 0.832 | 0.782 | - | 0.850 | 0.814 | 0.843 |
| EMD-9915 | 6K4M | 0.839 | 0.673 | - | 0.839 | 0.865 | 0.855 |
| EMD-20551 | 6Q0K | 0.613 | 0.576 | 0.595 | 0.565 | 0.606 | 0.626 |
| EMD-4975 | 6ROW | 0.739 | 0.739 | 0.695 | 0.718 | 0.700 | 0.718 |
| EMD-20724 | 6UC0 | 0.848 | 0.799 | 0.771 | 0.857 | 0.854 | 0.856 |
| EMD-21913 | 6WUH | 0.779 | 0.764 | 0.764 | 0.826 | 0.837 | 0.830 |
| EMD-11439 | 6ZU9 | 0.693 | 0.616 | 0.604 | 0.399 | 0.657 | 0.684 |
| EMD-12054 | 7B6H | 0.746 | 0.735 | 0.590 | 0.756 | 0.636 | 0.747 |
| EMD-30342 | 7CEC | 0.816 | 0.701 | 0.620 | 0.817 | 0.797 | 0.811 |
| EMD-30346 | 7CFS | 0.819 | 0.845 | 0.751 | 0.850 | 0.852 | 0.848 |
| EMD-30384 | 7CKB | 0.856 | 0.848 | - | 0.784 | 0.874 | 0.866 |
| EMD-30535 | 7D0I | 0.812 | 0.839 | 0.765 | 0.855 | 0.857 | 0.834 |
| EMD-31305 | 7EU0 | 0.766 | 0.776 | 0.732 | 0.799 | 0.788 | 0.783 |
| EMD-31561 | 7FER | 0.794 | 0.796 | 0.763 | 0.792 | 0.785 | 0.800 |
| EMD-31595 | 7FIF | 0.756 | 0.786 | 0.508 | 0.749 | 0.707 | 0.846 |
| EMD-22359 | 7JK2 | 0.861 | 0.771 | - | 0.819 | 0.826 | 0.830 |
| EMD-22375 | 7JLO | 0.836 | 0.860 | 0.810 | 0.873 | 0.870 | 0.869 |
| EMD-22445 | 7JRG | 0.876 | 0.880 | 0.837 | 0.860 | 0.878 | 0.881 |
| EMD-22233 | 7KEU | 0.742 | 0.784 | 0.583 | 0.815 | 0.714 | 0.752 |
| EMD-22879 | 7KHF | 0.833 | 0.853 | 0.820 | 0.870 | 0.882 | 0.879 |
| EMD-22901 | 7KJX | 0.803 | 0.809 | 0.749 | 0.802 | 0.839 | 0.841 |
| EMD-23340 | 7LHH | 0.743 | 0.639 | 0.596 | 0.643 | 0.755 | 0.749 |
| EMD-23526 | 7LUP | 0.856 | 0.832 | 0.795 | 0.844 | 0.857 | 0.858 |
| EMD-23930 | 7MP6 | 0.696 | 0.637 | 0.656 | 0.591 | 0.719 | 0.710 |
| EMD-24073 | 7MXL | 0.844 | 0.848 | - | 0.879 | 0.862 | 0.863 |
| EMD-12296 | 7NF6 | 0.851 | 0.811 | 0.782 | 0.879 | 0.867 | 0.856 |
| EMD-12309 | 7NG8 | 0.781 | 0.843 | 0.792 | 0.819 | 0.851 | 0.846 |
| EMD-12600 | 7NUR | 0.861 | 0.870 | 0.792 | 0.840 | 0.862 | 0.861 |
| EMD-12808 | 7OCI | 0.847 | 0.866 | 0.782 | 0.828 | 0.858 | 0.854 |
| EMD-12993 | 7ON1 | 0.823 | 0.836 | 0.716 | 0.785 | 0.822 | 0.817 |
| EMD-13035 | 7OQZ | 0.781 | 0.778 | 0.761 | 0.815 | 0.793 | 0.816 |
| EMD-13358 | 7PEV | 0.836 | 0.721 | 0.727 | 0.759 | 0.837 | 0.869 |
| EMD-13587 | 7PQ5 | 0.867 | 0.852 | 0.845 | 0.870 | 0.888 | 0.894 |

| | | | | | | | |
|---|---|---|---|---|---|---|---|
| EMD-13827 | 7Q56 | 0.895 | 0.823 | 0.755 | 0.888 | 0.890 | 0.881 |
| EMD-14145 | 7QTQ | 0.805 | 0.791 | - | 0.805 | 0.815 | 0.816 |
| EMD-14244 | 7R21 | 0.803 | 0.759 | 0.765 | 0.789 | 0.813 | 0.805 |
| EMD-24473 | 7RI4 | 0.759 | 0.628 | 0.736 | 0.800 | 0.790 | 0.793 |
| EMD-24889 | 7S82 | 0.803 | 0.716 | - | 0.775 | 0.845 | 0.850 |
| EMD-24991 | 7SBX | 0.907 | 0.903 | 0.876 | 0.869 | 0.886 | 0.891 |
| EMD-26377 | 7U6O | 0.784 | 0.533 | 0.737 | 0.851 | 0.795 | 0.815 |
| EMD-26512 | 7UHC | 0.796 | 0.758 | 0.757 | 0.826 | 0.830 | 0.827 |
| EMD-32536 | 7WIU | 0.845 | 0.745 | 0.796 | 0.834 | 0.833 | 0.856 |
| EMD-32845 | 7WV4 | 0.755 | 0.748 | - | 0.751 | 0.775 | 0.786 |
| EMD-32894 | 7WYU | 0.809 | 0.781 | 0.778 | 0.815 | 0.801 | 0.813 |
| EMD-33180 | 7XFZ | 0.856 | 0.868 | 0.787 | 0.807 | 0.846 | 0.833 |
| EMD-33192 | 7XHA | 0.758 | 0.787 | 0.745 | 0.818 | 0.823 | 0.817 |
| EMD-33666 | 7Y7I | 0.770 | 0.842 | 0.739 | 0.793 | 0.784 | 0.797 |
| EMD-33995 | 7YP7 | 0.826 | 0.849 | 0.762 | 0.856 | 0.846 | 0.856 |
| EMD-14849 | 7ZOX | 0.793 | 0.836 | 0.575 | 0.774 | 0.791 | 0.821 |
| EMD-15330 | 8AC1 | 0.777 | 0.662 | 0.734 | 0.771 | 0.774 | 0.774 |
| EMD-16555 | 8CC6 | 0.837 | 0.855 | 0.816 | 0.888 | 0.877 | 0.873 |
| EMD-26978 | 8CT2 | 0.863 | 0.787 | 0.830 | 0.775 | 0.878 | 0.871 |
| EMD-27034 | 8CX2 | 0.756 | 0.663 | 0.733 | 0.774 | 0.777 | 0.774 |
| EMD-27104 | 8D0B | 0.825 | 0.563 | 0.737 | 0.815 | 0.793 | 0.820 |
| EMD-27178 | 8D49 | 0.836 | 0.771 | 0.824 | 0.859 | 0.860 | 0.865 |
| EMD-27533 | 8DMB | 0.812 | 0.833 | 0.802 | 0.853 | 0.849 | 0.855 |
| EMD-28161 | 8EIQ | 0.871 | 0.885 | 0.839 | 0.811 | 0.869 | 0.880 |
| EMD-28204 | 8EKI | 0.761 | 0.547 | 0.625 | 0.753 | 0.748 | 0.792 |
| EMD-28534 | 8EPX | 0.833 | 0.786 | 0.823 | 0.870 | 0.862 | 0.870 |
| EMD-29686 | 8G30 | 0.872 | 0.742 | 0.824 | 0.860 | 0.858 | 0.871 |
| EMD-29936 | 8GCN | 0.823 | 0.495 | 0.697 | 0.725 | 0.798 | 0.839 |
| EMD-35304 | 8IAJ | 0.868 | 0.881 | 0.836 | 0.887 | 0.886 | 0.886 |
| EMD-35728 | 8IUO | 0.790 | 0.794 | 0.739 | 0.758 | 0.786 | 0.810 |
| EMD-35912 | 8J12 | 0.838 | 0.816 | 0.763 | 0.789 | 0.857 | 0.863 |
| EMD-35948 | 8J2F | 0.856 | 0.867 | 0.801 | 0.833 | 0.851 | 0.852 |
| EMD-36046 | 8J7R | 0.765 | 0.706 | 0.664 | 0.768 | 0.738 | 0.806 |
| EMD-36070 | 8J8H | 0.773 | 0.568 | 0.693 | 0.783 | 0.754 | 0.794 |
| EMD-36824 | 8K21 | 0.758 | 0.790 | 0.724 | 0.807 | 0.814 | 0.813 |
| EMD-17341 | 8P0X | 0.796 | 0.762 | 0.694 | 0.700 | 0.755 | 0.784 |
| EMD-17632 | 8PEN | 0.871 | 0.642 | 0.776 | 0.791 | 0.810 | 0.836 |
| EMD-17979 | 8PVX | 0.866 | 0.777 | 0.768 | 0.842 | 0.811 | 0.844 |
| EMD-18345 | 8QE8 | 0.800 | 0.850 | 0.735 | 0.816 | 0.863 | 0.863 |
| EMD-18453 | 8QJX | 0.865 | 0.849 | 0.842 | 0.897 | 0.867 | 0.882 |
| EMD-18471 | 8QKU | 0.844 | 0.746 | 0.678 | 0.829 | 0.790 | 0.827 |
| EMD-18608 | 8QQX | 0.877 | 0.889 | 0.832 | 0.876 | 0.882 | 0.881 |
| EMD-18638 | 8QSP | 0.847 | 0.508 | 0.719 | 0.775 | 0.775 | 0.830 |

| | | | | | | | |
|---|---|---|---|---|---|---|---|
| EMD-18778 | 8QZM | 0.872 | 0.795 | 0.853 | 0.859 | 0.877 | 0.855 |
| EMD-40241 | 8S9P | 0.769 | 0.774 | 0.696 | 0.763 | 0.799 | 0.800 |
| EMD-41011 | 8T3T | 0.909 | 0.896 | 0.848 | 0.842 | 0.853 | 0.868 |
| EMD-41355 | 8TKO | 0.857 | 0.704 | 0.774 | 0.813 | 0.804 | 0.827 |
| EMD-42774 | 8UXQ | 0.633 | 0.611 | 0.580 | 0.558 | 0.604 | 0.641 |
| EMD-42795 | 8UY8 | 0.880 | 0.905 | 0.804 | 0.842 | 0.865 | 0.869 |
| EMD-42960 | 8V3Y | 0.844 | 0.811 | - | 0.837 | 0.842 | 0.845 |
| EMD-43196 | 8VG0 | 0.817 | 0.610 | 0.748 | 0.838 | 0.797 | 0.831 |
| EMD-43304 | 8VK4 | 0.863 | 0.860 | 0.739 | 0.798 | 0.806 | 0.818 |
| EMD-43316 | 8VKH | 0.833 | 0.794 | 0.761 | 0.859 | 0.804 | 0.835 |
| EMD-43346 | 8VLW | 0.871 | 0.734 | 0.814 | 0.835 | 0.848 | 0.874 |
| EMD-37439 | 8WCE | 0.818 | 0.814 | 0.774 | 0.804 | 0.841 | 0.831 |
| EMD-37658 | 8WMQ | 0.803 | 0.806 | 0.771 | 0.789 | 0.808 | 0.833 |
| EMD-38025 | 8X31 | 0.649 | 0.594 | 0.444 | 0.616 | 0.555 | 0.716 |
| EMD-39454 | 8YON | 0.751 | 0.669 | 0.600 | 0.726 | 0.656 | 0.720 |
| EMD-60471 | 8ZTS | 0.791 | 0.862 | 0.755 | 0.784 | 0.835 | 0.861 |
| EMD-44441 | 9BCX | 0.658 | 0.593 | 0.624 | 0.637 | 0.671 | 0.679 |
| EMD-45805 | 9CPO | 0.756 | 0.743 | 0.767 | 0.839 | 0.824 | 0.833 |
| EMD-45924 | 9CU0 | 0.792 | 0.706 | 0.737 | 0.785 | 0.784 | 0.820 |
| EMD-45988 | 9CXF | 0.902 | 0.752 | 0.779 | 0.849 | 0.850 | 0.905 |
| EMD-46645 | 9D8P | 0.842 | 0.740 | 0.797 | 0.890 | 0.830 | 0.854 |
| EMD-47091 | 9DOU | 0.827 | 0.864 | 0.763 | 0.822 | 0.839 | 0.845 |
| EMD-47476 | 9E3A | 0.801 | 0.656 | 0.756 | 0.830 | 0.805 | 0.820 |
| EMD-50262 | 9F9X | 0.880 | 0.770 | - | 0.800 | 0.799 | 0.830 |
| EMD-50289 | 9FB6 | 0.886 | 0.852 | 0.826 | 0.809 | 0.847 | 0.854 |
| EMD-50522 | 9FKD | 0.855 | 0.720 | 0.783 | 0.807 | 0.836 | 0.855 |
| EMD-51241 | 9GD1 | 0.863 | 0.703 | 0.800 | 0.822 | 0.817 | 0.841 |
| EMD-51918 | 9H82 | 0.768 | 0.688 | 0.569 | 0.439 | 0.698 | 0.695 |
| EMD-61073 | 9J1J | 0.853 | 0.624 | 0.759 | 0.847 | 0.774 | 0.812 |
| EMD-62812 | 9L4F | 0.769 | 0.696 | 0.761 | 0.689 | 0.780 | 0.815 |

**Supplementary Table 5 | Summary of CC_box for each case in the primary-map dataset.** A dash ("-") indicates that the method failed to process the case.

| CC_box | | | | | | | |
|---|---|---|---|---|---|---|---|
| EMDBID | PDBID | Deposited | phenix | DeepEMhancer | CryoTEN | EMReady | DEMO-EMReF |
| EMD-3605 | 5N9Y | 0.782 | 0.725 | 0.630 | 0.937 | 0.898 | 0.889 |
| EMD-8745 | 5VY9 | 0.841 | 0.710 | 0.663 | 0.868 | 0.867 | 0.876 |
| EMD-6714 | 5XB1 | 0.799 | 0.887 | 0.880 | 0.915 | 0.920 | 0.903 |
| EMD-7065 | 6B7Y | 0.885 | 0.742 | 0.632 | 0.926 | 0.914 | 0.948 |
| EMD-3896 | 6EMK | 0.786 | 0.672 | 0.668 | 0.801 | 0.799 | 0.821 |
| EMD-0088 | 6GY6 | 0.894 | 0.760 | - | 0.925 | 0.888 | 0.902 |
| EMD-9915 | 6K4M | 0.678 | 0.463 | - | 0.806 | 0.833 | 0.869 |
| EMD-20551 | 6Q0K | 0.741 | 0.536 | 0.687 | 0.758 | 0.791 | 0.811 |
| EMD-4975 | 6ROW | 0.808 | 0.787 | 0.718 | 0.821 | 0.817 | 0.833 |
| EMD-20724 | 6UC0 | 0.783 | 0.736 | 0.673 | 0.851 | 0.817 | 0.806 |
| EMD-21913 | 6WUH | 0.666 | 0.613 | 0.770 | 0.893 | 0.900 | 0.889 |
| EMD-11439 | 6ZU9 | 0.861 | 0.707 | 0.788 | 0.774 | 0.881 | 0.897 |
| EMD-12054 | 7B6H | 0.796 | 0.784 | 0.644 | 0.877 | 0.806 | 0.875 |
| EMD-30342 | 7CEC | 0.902 | 0.775 | 0.633 | 0.920 | 0.903 | 0.905 |
| EMD-30346 | 7CFS | 0.784 | 0.796 | 0.764 | 0.914 | 0.909 | 0.899 |
| EMD-30384 | 7CKB | 0.835 | 0.862 | - | 0.839 | 0.903 | 0.897 |
| EMD-30535 | 7D0I | 0.689 | 0.757 | 0.717 | 0.901 | 0.901 | 0.882 |
| EMD-31305 | 7EU0 | 0.728 | 0.815 | 0.717 | 0.852 | 0.852 | 0.848 |
| EMD-31561 | 7FER | 0.665 | 0.643 | 0.734 | 0.836 | 0.842 | 0.845 |
| EMD-31595 | 7FIF | 0.650 | 0.595 | 0.489 | 0.855 | 0.853 | 0.924 |
| EMD-22359 | 7JK2 | 0.848 | 0.618 | - | 0.852 | 0.870 | 0.865 |
| EMD-22375 | 7JLO | 0.701 | 0.746 | 0.827 | 0.913 | 0.910 | 0.909 |
| EMD-22445 | 7JRG | 0.787 | 0.790 | 0.846 | 0.906 | 0.925 | 0.925 |
| EMD-22233 | 7KEU | 0.821 | 0.785 | 0.630 | 0.923 | 0.846 | 0.859 |
| EMD-22879 | 7KHF | 0.777 | 0.875 | 0.825 | 0.930 | 0.934 | 0.929 |
| EMD-22901 | 7KJX | 0.710 | 0.732 | 0.764 | 0.865 | 0.896 | 0.895 |
| EMD-23340 | 7LHH | 0.717 | 0.567 | 0.615 | 0.840 | 0.903 | 0.909 |
| EMD-23526 | 7LUP | 0.864 | 0.772 | 0.797 | 0.938 | 0.932 | 0.937 |
| EMD-23930 | 7MP6 | 0.848 | 0.657 | 0.724 | 0.874 | 0.854 | 0.878 |
| EMD-24073 | 7MXL | 0.782 | 0.787 | - | 0.935 | 0.917 | 0.909 |
| EMD-12296 | 7NF6 | 0.692 | 0.612 | 0.713 | 0.878 | 0.881 | 0.871 |
| EMD-12309 | 7NG8 | 0.868 | 0.813 | 0.829 | 0.883 | 0.904 | 0.902 |
| EMD-12600 | 7NUR | 0.766 | 0.877 | 0.778 | 0.887 | 0.904 | 0.901 |
| EMD-12808 | 7OCI | 0.643 | 0.707 | 0.724 | 0.879 | 0.908 | 0.900 |
| EMD-12993 | 7ON1 | 0.813 | 0.869 | 0.687 | 0.877 | 0.895 | 0.895 |
| EMD-13035 | 7OQZ | 0.611 | 0.607 | 0.753 | 0.862 | 0.847 | 0.859 |
| EMD-13358 | 7PEV | 0.908 | 0.788 | 0.746 | 0.875 | 0.910 | 0.951 |
| EMD-13587 | 7PQ5 | 0.847 | 0.760 | 0.859 | 0.895 | 0.917 | 0.920 |

| | | | | | | | |
|---|---|---|---|---|---|---|---|
| EMD-13827 | 7Q56 | 0.934 | 0.829 | 0.782 | 0.961 | 0.954 | 0.959 |
| EMD-14145 | 7QTQ | 0.716 | 0.620 | - | 0.833 | 0.829 | 0.848 |
| EMD-14244 | 7R21 | 0.736 | 0.629 | 0.796 | 0.878 | 0.880 | 0.872 |
| EMD-24473 | 7RI4 | 0.703 | 0.439 | 0.747 | 0.889 | 0.874 | 0.866 |
| EMD-24889 | 7S82 | 0.775 | 0.508 | - | 0.830 | 0.901 | 0.899 |
| EMD-24991 | 7SBX | 0.855 | 0.867 | 0.887 | 0.915 | 0.924 | 0.929 |
| EMD-26377 | 7U6O | 0.854 | 0.459 | 0.738 | 0.928 | 0.856 | 0.861 |
| EMD-26512 | 7UHC | 0.781 | 0.618 | 0.801 | 0.867 | 0.873 | 0.881 |
| EMD-32536 | 7WIU | 0.675 | 0.501 | 0.688 | 0.878 | 0.890 | 0.892 |
| EMD-32845 | 7WV4 | 0.791 | 0.757 | - | 0.844 | 0.858 | 0.878 |
| EMD-32894 | 7WYU | 0.738 | 0.654 | 0.804 | 0.895 | 0.879 | 0.883 |
| EMD-33180 | 7XFZ | 0.774 | 0.839 | 0.798 | 0.874 | 0.900 | 0.887 |
| EMD-33192 | 7XHA | 0.686 | 0.722 | 0.759 | 0.892 | 0.889 | 0.880 |
| EMD-33666 | 7Y7I | 0.645 | 0.840 | 0.773 | 0.874 | 0.866 | 0.877 |
| EMD-33995 | 7YP7 | 0.782 | 0.843 | 0.711 | 0.897 | 0.887 | 0.891 |
| EMD-14849 | 7ZOX | 0.849 | 0.831 | 0.634 | 0.896 | 0.892 | 0.895 |
| EMD-15330 | 8AC1 | 0.826 | 0.821 | 0.764 | 0.880 | 0.873 | 0.878 |
| EMD-16555 | 8CC6 | 0.700 | 0.741 | 0.827 | 0.925 | 0.917 | 0.911 |
| EMD-26978 | 8CT2 | 0.617 | 0.528 | 0.806 | 0.778 | 0.889 | 0.874 |
| EMD-27034 | 8CX2 | 0.831 | 0.428 | 0.753 | 0.864 | 0.864 | 0.867 |
| EMD-27104 | 8D0B | 0.872 | 0.366 | 0.729 | 0.878 | 0.856 | 0.865 |
| EMD-27178 | 8D49 | 0.855 | 0.638 | 0.837 | 0.896 | 0.892 | 0.901 |
| EMD-27533 | 8DMB | 0.688 | 0.826 | 0.826 | 0.912 | 0.905 | 0.911 |
| EMD-28161 | 8EIQ | 0.651 | 0.692 | 0.819 | 0.827 | 0.896 | 0.900 |
| EMD-28204 | 8EKI | 0.858 | 0.545 | 0.629 | 0.892 | 0.868 | 0.862 |
| EMD-28534 | 8EPX | 0.789 | 0.675 | 0.847 | 0.916 | 0.915 | 0.917 |
| EMD-29686 | 8G30 | 0.828 | 0.550 | 0.758 | 0.808 | 0.825 | 0.836 |
| EMD-29936 | 8GCN | 0.899 | 0.474 | 0.742 | 0.873 | 0.894 | 0.907 |
| EMD-35304 | 8IAJ | 0.735 | 0.784 | 0.857 | 0.923 | 0.923 | 0.923 |
| EMD-35728 | 8IUO | 0.605 | 0.582 | 0.601 | 0.757 | 0.790 | 0.778 |
| EMD-35912 | 8J12 | 0.843 | 0.755 | 0.759 | 0.907 | 0.910 | 0.907 |
| EMD-35948 | 8J2F | 0.661 | 0.710 | 0.797 | 0.879 | 0.901 | 0.896 |
| EMD-36046 | 8J7R | 0.690 | 0.496 | 0.659 | 0.813 | 0.818 | 0.850 |
| EMD-36070 | 8J8H | 0.823 | 0.497 | 0.709 | 0.868 | 0.832 | 0.855 |
| EMD-36824 | 8K21 | 0.775 | 0.715 | 0.615 | 0.767 | 0.784 | 0.782 |
| EMD-17341 | 8P0X | 0.841 | 0.750 | 0.768 | 0.862 | 0.912 | 0.922 |
| EMD-17632 | 8PEN | 0.908 | 0.454 | 0.744 | 0.846 | 0.867 | 0.881 |
| EMD-17979 | 8PVX | 0.779 | 0.701 | 0.633 | 0.742 | 0.732 | 0.738 |
| EMD-18345 | 8QE8 | 0.780 | 0.845 | 0.736 | 0.834 | 0.905 | 0.901 |
| EMD-18453 | 8QJX | 0.879 | 0.820 | 0.832 | 0.948 | 0.909 | 0.920 |
| EMD-18471 | 8QKU | 0.913 | 0.759 | 0.661 | 0.912 | 0.873 | 0.899 |
| EMD-18608 | 8QQX | 0.912 | 0.823 | 0.831 | 0.918 | 0.924 | 0.925 |
| EMD-18638 | 8QSP | 0.912 | 0.381 | 0.750 | 0.878 | 0.870 | 0.901 |

| | | | | | | | |
|---|---|---|---|---|---|---|---|
| EMD-18778 | 8QZM | 0.905 | 0.631 | 0.854 | 0.914 | 0.924 | 0.923 |
| EMD-40241 | 8S9P | 0.796 | 0.777 | 0.729 | 0.862 | 0.880 | 0.875 |
| EMD-41011 | 8T3T | 0.884 | 0.824 | 0.846 | 0.899 | 0.902 | 0.920 |
| EMD-41355 | 8TKO | 0.877 | 0.604 | 0.766 | 0.872 | 0.869 | 0.877 |
| EMD-42774 | 8UXQ | 0.813 | 0.612 | 0.653 | 0.730 | 0.762 | 0.807 |
| EMD-42795 | 8UY8 | 0.685 | 0.737 | 0.756 | 0.879 | 0.904 | 0.900 |
| EMD-42960 | 8V3Y | 0.670 | 0.615 | - | 0.675 | 0.772 | 0.802 |
| EMD-43196 | 8VG0 | 0.799 | 0.457 | 0.753 | 0.894 | 0.856 | 0.884 |
| EMD-43304 | 8VK4 | 0.898 | 0.884 | 0.697 | 0.848 | 0.876 | 0.871 |
| EMD-43316 | 8VKH | 0.790 | 0.715 | 0.721 | 0.872 | 0.826 | 0.840 |
| EMD-43346 | 8VLW | 0.807 | 0.515 | 0.796 | 0.875 | 0.895 | 0.912 |
| EMD-37439 | 8WCE | 0.748 | 0.650 | 0.820 | 0.870 | 0.889 | 0.881 |
| EMD-37658 | 8WMQ | 0.621 | 0.600 | 0.768 | 0.834 | 0.867 | 0.876 |
| EMD-38025 | 8X31 | 0.736 | 0.516 | 0.491 | 0.760 | 0.661 | 0.861 |
| EMD-39454 | 8YON | 0.846 | 0.726 | 0.720 | 0.877 | 0.798 | 0.900 |
| EMD-60471 | 8ZTS | 0.631 | 0.766 | 0.708 | 0.843 | 0.875 | 0.901 |
| EMD-44441 | 9BCX | 0.804 | 0.560 | 0.699 | 0.823 | 0.831 | 0.845 |
| EMD-45805 | 9CPO | 0.652 | 0.591 | 0.780 | 0.904 | 0.887 | 0.881 |
| EMD-45924 | 9CU0 | 0.778 | 0.498 | 0.745 | 0.877 | 0.864 | 0.880 |
| EMD-45988 | 9CXF | 0.925 | 0.533 | 0.760 | 0.902 | 0.907 | 0.941 |
| EMD-46645 | 9D8P | 0.797 | 0.599 | 0.743 | 0.890 | 0.837 | 0.837 |
| EMD-47091 | 9DOU | 0.632 | 0.779 | 0.767 | 0.872 | 0.890 | 0.892 |
| EMD-47476 | 9E3A | 0.687 | 0.441 | 0.603 | 0.732 | 0.714 | 0.730 |
| EMD-50262 | 9F9X | 0.933 | 0.756 | - | 0.865 | 0.863 | 0.888 |
| EMD-50289 | 9FB6 | 0.921 | 0.746 | 0.813 | 0.880 | 0.902 | 0.900 |
| EMD-50522 | 9FKD | 0.853 | 0.500 | 0.790 | 0.853 | 0.891 | 0.895 |
| EMD-51241 | 9GD1 | 0.915 | 0.699 | 0.771 | 0.902 | 0.907 | 0.926 |
| EMD-51918 | 9H82 | 0.846 | 0.514 | 0.708 | 0.623 | 0.843 | 0.834 |
| EMD-61073 | 9J1J | 0.837 | 0.520 | 0.714 | 0.856 | 0.797 | 0.828 |
| EMD-62812 | 9L4F | 0.785 | 0.590 | 0.807 | 0.831 | 0.871 | 0.888 |

**Supplementary Table 6 | Summary of CC_peaks for each case in the primary-map dataset.** A dash ("-") indicates that the method failed to process the case.

| CC_peaks | | | | | | | |
|---|---|---|---|---|---|---|---|
| EMDBID | PDBID | Deposited | phenix | DeepEMhancer | CryoTEN | EMReady | DEMO-EMReF |
| EMD-3605 | 5N9Y | 0.649 | 0.601 | 0.536 | 0.837 | 0.819 | 0.827 |
| EMD-8745 | 5VY9 | 0.675 | 0.591 | 0.583 | 0.695 | 0.712 | 0.718 |
| EMD-6714 | 5XB1 | 0.795 | 0.860 | 0.874 | 0.893 | 0.900 | 0.878 |
| EMD-7065 | 6B7Y | 0.693 | 0.433 | 0.429 | 0.727 | 0.662 | 0.770 |
| EMD-3896 | 6EMK | 0.473 | 0.345 | 0.403 | 0.381 | 0.335 | 0.418 |
| EMD-0088 | 6GY6 | 0.795 | 0.682 | - | 0.837 | 0.793 | 0.839 |
| EMD-9915 | 6K4M | 0.359 | 0.260 | - | 0.546 | 0.613 | 0.778 |
| EMD-20551 | 6Q0K | 0.463 | 0.270 | 0.463 | 0.382 | 0.428 | 0.490 |
| EMD-4975 | 6ROW | 0.640 | 0.617 | 0.652 | 0.606 | 0.603 | 0.632 |
| EMD-20724 | 6UC0 | 0.547 | 0.491 | 0.480 | 0.599 | 0.503 | 0.528 |
| EMD-21913 | 6WUH | 0.629 | 0.597 | 0.703 | 0.821 | 0.836 | 0.830 |
| EMD-11439 | 6ZU9 | 0.644 | 0.527 | 0.575 | 0.361 | 0.609 | 0.646 |
| EMD-12054 | 7B6H | 0.613 | 0.579 | 0.529 | 0.653 | 0.480 | 0.623 |
| EMD-30342 | 7CEC | 0.760 | 0.573 | 0.507 | 0.780 | 0.757 | 0.777 |
| EMD-30346 | 7CFS | 0.771 | 0.786 | 0.717 | 0.853 | 0.858 | 0.855 |
| EMD-30384 | 7CKB | 0.786 | 0.820 | - | 0.786 | 0.880 | 0.873 |
| EMD-30535 | 7D0I | 0.647 | 0.715 | 0.693 | 0.854 | 0.866 | 0.843 |
| EMD-31305 | 7EU0 | 0.671 | 0.746 | 0.686 | 0.794 | 0.798 | 0.797 |
| EMD-31561 | 7FER | 0.547 | 0.526 | 0.695 | 0.709 | 0.749 | 0.759 |
| EMD-31595 | 7FIF | 0.485 | 0.504 | 0.222 | 0.660 | 0.678 | 0.821 |
| EMD-22359 | 7JK2 | 0.736 | 0.516 | - | 0.764 | 0.811 | 0.824 |
| EMD-22375 | 7JLO | 0.676 | 0.714 | 0.813 | 0.877 | 0.874 | 0.876 |
| EMD-22445 | 7JRG | 0.747 | 0.747 | 0.822 | 0.849 | 0.882 | 0.886 |
| EMD-22233 | 7KEU | 0.742 | 0.700 | 0.555 | 0.815 | 0.706 | 0.747 |
| EMD-22879 | 7KHF | 0.754 | 0.838 | 0.791 | 0.879 | 0.889 | 0.886 |
| EMD-22901 | 7KJX | 0.660 | 0.676 | 0.729 | 0.793 | 0.844 | 0.848 |
| EMD-23340 | 7LHH | 0.461 | 0.324 | 0.384 | 0.574 | 0.736 | 0.717 |
| EMD-23526 | 7LUP | 0.796 | 0.685 | 0.768 | 0.833 | 0.845 | 0.840 |
| EMD-23930 | 7MP6 | 0.629 | 0.461 | 0.512 | 0.576 | 0.587 | 0.632 |
| EMD-24073 | 7MXL | 0.762 | 0.765 | - | 0.876 | 0.860 | 0.862 |
| EMD-12296 | 7NF6 | 0.650 | 0.590 | 0.682 | 0.806 | 0.818 | 0.810 |
| EMD-12309 | 7NG8 | 0.785 | 0.765 | 0.785 | 0.820 | 0.863 | 0.861 |
| EMD-12600 | 7NUR | 0.736 | 0.813 | 0.702 | 0.817 | 0.857 | 0.858 |
| EMD-12808 | 7OCI | 0.660 | 0.698 | 0.686 | 0.811 | 0.857 | 0.857 |
| EMD-12993 | 7ON1 | 0.715 | 0.769 | 0.596 | 0.773 | 0.829 | 0.829 |
| EMD-13035 | 7OQZ | 0.527 | 0.528 | 0.732 | 0.794 | 0.781 | 0.807 |
| EMD-13358 | 7PEV | 0.774 | 0.601 | 0.621 | 0.741 | 0.823 | 0.830 |
| EMD-13587 | 7PQ5 | 0.816 | 0.742 | 0.839 | 0.847 | 0.874 | 0.881 |

| | | | | | | | |
|---|---|---|---|---|---|---|---|
| EMD-13827 | 7Q56 | 0.884 | 0.732 | 0.696 | 0.886 | 0.890 | 0.878 |
| EMD-14145 | 7QTQ | 0.615 | 0.563 | - | 0.701 | 0.721 | 0.763 |
| EMD-14244 | 7R21 | 0.706 | 0.599 | 0.777 | 0.808 | 0.822 | 0.821 |
| EMD-24473 | 7RI4 | 0.603 | 0.379 | 0.689 | 0.797 | 0.790 | 0.796 |
| EMD-24889 | 7S82 | 0.709 | 0.442 | - | 0.755 | 0.842 | 0.849 |
| EMD-24991 | 7SBX | 0.856 | 0.859 | 0.870 | 0.873 | 0.885 | 0.893 |
| EMD-26377 | 7U6O | 0.692 | 0.370 | 0.711 | 0.833 | 0.778 | 0.807 |
| EMD-26512 | 7UHC | 0.738 | 0.586 | 0.759 | 0.805 | 0.810 | 0.823 |
| EMD-32536 | 7WIU | 0.549 | 0.370 | 0.644 | 0.788 | 0.821 | 0.829 |
| EMD-32845 | 7WV4 | 0.610 | 0.577 | - | 0.643 | 0.716 | 0.734 |
| EMD-32894 | 7WYU | 0.677 | 0.588 | 0.756 | 0.824 | 0.813 | 0.826 |
| EMD-33180 | 7XFZ | 0.764 | 0.811 | 0.744 | 0.810 | 0.858 | 0.847 |
| EMD-33192 | 7XHA | 0.631 | 0.660 | 0.703 | 0.831 | 0.835 | 0.830 |
| EMD-33666 | 7Y7I | 0.591 | 0.785 | 0.747 | 0.797 | 0.794 | 0.809 |
| EMD-33995 | 7YP7 | 0.716 | 0.771 | 0.641 | 0.837 | 0.827 | 0.847 |
| EMD-14849 | 7ZOX | 0.712 | 0.727 | 0.497 | 0.749 | 0.785 | 0.820 |
| EMD-15330 | 8AC1 | 0.666 | 0.582 | 0.649 | 0.714 | 0.751 | 0.777 |
| EMD-16555 | 8CC6 | 0.691 | 0.730 | 0.815 | 0.889 | 0.881 | 0.879 |
| EMD-26978 | 8CT2 | 0.637 | 0.525 | 0.760 | 0.700 | 0.842 | 0.823 |
| EMD-27034 | 8CX2 | 0.688 | 0.321 | 0.670 | 0.735 | 0.774 | 0.772 |
| EMD-27104 | 8D0B | 0.774 | 0.266 | 0.674 | 0.795 | 0.782 | 0.814 |
| EMD-27178 | 8D49 | 0.796 | 0.611 | 0.806 | 0.836 | 0.831 | 0.845 |
| EMD-27533 | 8DMB | 0.670 | 0.789 | 0.805 | 0.856 | 0.854 | 0.859 |
| EMD-28161 | 8EIQ | 0.613 | 0.665 | 0.789 | 0.765 | 0.843 | 0.852 |
| EMD-28204 | 8EKI | 0.696 | 0.338 | 0.543 | 0.732 | 0.725 | 0.777 |
| EMD-28534 | 8EPX | 0.760 | 0.649 | 0.818 | 0.868 | 0.869 | 0.873 |
| EMD-29686 | 8G30 | 0.693 | 0.504 | 0.674 | 0.621 | 0.661 | 0.677 |
| EMD-29936 | 8GCN | 0.739 | 0.309 | 0.613 | 0.681 | 0.780 | 0.826 |
| EMD-35304 | 8IAJ | 0.723 | 0.772 | 0.843 | 0.890 | 0.892 | 0.893 |
| EMD-35728 | 8IUO | 0.396 | 0.370 | 0.386 | 0.454 | 0.565 | 0.550 |
| EMD-35912 | 8J12 | 0.824 | 0.750 | 0.743 | 0.800 | 0.857 | 0.866 |
| EMD-35948 | 8J2F | 0.632 | 0.672 | 0.750 | 0.815 | 0.852 | 0.857 |
| EMD-36046 | 8J7R | 0.552 | 0.632 | 0.595 | 0.595 | 0.664 | 0.688 |
| EMD-36070 | 8J8H | 0.699 | 0.402 | 0.635 | 0.755 | 0.738 | 0.776 |
| EMD-36824 | 8K21 | 0.562 | 0.541 | 0.494 | 0.534 | 0.578 | 0.575 |
| EMD-17341 | 8P0X | 0.683 | 0.623 | 0.640 | 0.576 | 0.631 | 0.661 |
| EMD-17632 | 8PEN | 0.812 | 0.404 | 0.657 | 0.766 | 0.806 | 0.835 |
| EMD-17979 | 8PVX | 0.522 | 0.468 | 0.452 | 0.478 | 0.466 | 0.481 |
| EMD-18345 | 8QE8 | 0.733 | 0.814 | 0.684 | 0.796 | 0.868 | 0.868 |
| EMD-18453 | 8QJX | 0.811 | 0.757 | 0.801 | 0.896 | 0.857 | 0.878 |
| EMD-18471 | 8QKU | 0.792 | 0.619 | 0.567 | 0.796 | 0.754 | 0.811 |
| EMD-18608 | 8QQX | 0.861 | 0.793 | 0.802 | 0.871 | 0.886 | 0.888 |
| EMD-18638 | 8QSP | 0.764 | 0.236 | 0.646 | 0.719 | 0.752 | 0.815 |

| | | | | | | | |
|---|---|---|---|---|---|---|---|
| EMD-18778 | 8QZM | 0.834 | 0.581 | 0.803 | 0.836 | 0.875 | 0.857 |
| EMD-40241 | 8S9P | 0.642 | 0.623 | 0.600 | 0.688 | 0.778 | 0.789 |
| EMD-41011 | 8T3T | 0.844 | 0.790 | 0.821 | 0.808 | 0.842 | 0.869 |
| EMD-41355 | 8TKO | 0.803 | 0.524 | 0.720 | 0.805 | 0.799 | 0.826 |
| EMD-42774 | 8UXQ | 0.551 | 0.438 | 0.508 | 0.517 | 0.573 | 0.610 |
| EMD-42795 | 8UY8 | 0.651 | 0.695 | 0.686 | 0.825 | 0.863 | 0.868 |
| EMD-42960 | 8V3Y | 0.548 | 0.507 | - | 0.407 | 0.588 | 0.653 |
| EMD-43196 | 8VG0 | 0.731 | 0.388 | 0.725 | 0.831 | 0.794 | 0.833 |
| EMD-43304 | 8VK4 | 0.757 | 0.744 | 0.629 | 0.760 | 0.800 | 0.816 |
| EMD-43316 | 8VKH | 0.621 | 0.553 | 0.596 | 0.662 | 0.610 | 0.630 |
| EMD-43346 | 8VLW | 0.705 | 0.471 | 0.730 | 0.813 | 0.848 | 0.875 |
| EMD-37439 | 8WCE | 0.724 | 0.618 | 0.791 | 0.811 | 0.852 | 0.844 |
| EMD-37658 | 8WMQ | 0.535 | 0.530 | 0.710 | 0.730 | 0.797 | 0.825 |
| EMD-38025 | 8X31 | 0.571 | 0.419 | 0.397 | 0.537 | 0.484 | 0.654 |
| EMD-39454 | 8YON | 0.671 | 0.538 | 0.567 | 0.682 | 0.631 | 0.660 |
| EMD-60471 | 8ZTS | 0.523 | 0.664 | 0.675 | 0.767 | 0.818 | 0.857 |
| EMD-44441 | 9BCX | 0.560 | 0.361 | 0.557 | 0.568 | 0.614 | 0.605 |
| EMD-45805 | 9CPO | 0.622 | 0.570 | 0.755 | 0.841 | 0.827 | 0.838 |
| EMD-45924 | 9CU0 | 0.666 | 0.306 | 0.659 | 0.754 | 0.781 | 0.820 |
| EMD-45988 | 9CXF | 0.854 | 0.584 | 0.732 | 0.796 | 0.843 | 0.900 |
| EMD-46645 | 9D8P | 0.689 | 0.534 | 0.642 | 0.752 | 0.698 | 0.686 |
| EMD-47091 | 9DOU | 0.605 | 0.801 | 0.763 | 0.825 | 0.846 | 0.851 |
| EMD-47476 | 9E3A | 0.487 | 0.331 | 0.457 | 0.413 | 0.484 | 0.518 |
| EMD-50262 | 9F9X | 0.874 | 0.723 | - | 0.795 | 0.795 | 0.826 |
| EMD-50289 | 9FB6 | 0.876 | 0.744 | 0.797 | 0.803 | 0.850 | 0.854 |
| EMD-50522 | 9FKD | 0.779 | 0.485 | 0.745 | 0.782 | 0.833 | 0.855 |
| EMD-51241 | 9GD1 | 0.792 | 0.509 | 0.602 | 0.768 | 0.805 | 0.833 |
| EMD-51918 | 9H82 | 0.582 | 0.792 | 0.217 | -0.201 | 0.454 | 0.516 |
| EMD-61073 | 9J1J | 0.678 | 0.346 | 0.608 | 0.673 | 0.594 | 0.637 |
| EMD-62812 | 9L4F | 0.694 | 0.490 | 0.684 | 0.641 | 0.729 | 0.768 |

**Supplementary Table 7 | Average results of different methods on the 27 cryo-EM half-maps.** Bold values indicate the best performance for each metric.

| Methods | FSC-0.5 (Å) | Q-score | CC_mask | CC_box | CC_peaks |
|---|---|---|---|---|---|
| Averaged half-maps | 4.99 | 0.503 | 0.785 | 0.701 | 0.657 |
| phenix | 4.90 | 0.531 | 0.796 | 0.734 | 0.660 |
| DeepEMhancer | 4.01 | 0.468 | 0.782 | 0.798 | 0.749 |
| CryoTEN | 4.46 | 0.527 | 0.787 | 0.858 | 0.762 |
| EMReady | 3.64 | 0.569 | 0.811 | 0.880 | 0.791 |
| **DEMO-EMReF** | **3.44** | **0.571** | **0.819** | **0.886** | **0.798** |

**Supplementary Table 8 | Summary of FSC-0.5 resolution for the 27 half-maps processed by different methods.**

| FSC-0.5 | | | | | | | |
|---|---|---|---|---|---|---|---|
| EMDBID | PDBID | Averaged | phenix | DeepEMhancer | CryoTEN | EMReady | DEMO-EMReF |
| EMD-30535 | 7D0I | 4.38 | 4.38 | 3.07 | 2.98 | 2.32 | 2.21 |
| EMD-31595 | 7FIF | 8.01 | 7.67 | 6.14 | 6.55 | 7.65 | 6.77 |
| EMD-22375 | 7JLO | 3.82 | 3.81 | 3.24 | 3.08 | 2.23 | 2.04 |
| EMD-12296 | 7NF6 | 3.7 | 3.67 | 3.32 | 3.06 | 2.62 | 2.56 |
| EMD-13587 | 7PQ5 | 3.62 | 3.62 | 3.18 | 2.99 | 2.32 | 2.08 |
| EMD-26377 | 7U6O | 3.95 | 3.95 | 3.80 | 3.63 | 3.42 | 3.29 |
| EMD-33995 | 7YP7 | 4.36 | 4.38 | 3.37 | 3.23 | 3.05 | 2.91 |
| EMD-16555 | 8CC6 | 3.71 | 3.71 | 3.21 | 3.08 | 2.41 | 2.36 |
| EMD-27533 | 8DMB | 3.79 | 3.9 | 3.05 | 3 | 2.99 | 2.61 |
| EMD-29686 | 8G30 | 3.83 | 3.71 | 3.20 | 3.18 | 2.98 | 2.87 |
| EMD-29936 | 8GCN | 4.45 | 4.49 | 4.53 | 4.39 | 4.33 | 4.32 |
| EMD-35304 | 8IAJ | 3.61 | 3.61 | 3.18 | 2.97 | 2.28 | 2.09 |
| EMD-35912 | 8J12 | 3.57 | 3.57 | 2.98 | 3.03 | 2.61 | 2.06 |
| EMD-35948 | 8J2F | 3.46 | 3.46 | 3.25 | 3.13 | 2.62 | 2.25 |
| EMD-18345 | 8QE8 | 8.16 | 8.15 | 3.35 | 3.56 | 2.86 | 2.6 |
| EMD-18453 | 8QJX | 3.51 | 3.53 | 2.93 | 2.89 | 2.17 | 2.04 |
| EMD-18608 | 8QQX | 3.61 | 3.61 | 2.94 | 2.66 | 2.18 | 2.01 |
| EMD-18778 | 8QZM | 3.58 | 3.58 | 3.08 | 3 | 2.31 | 2.24 |
| EMD-41355 | 8TKO | 4.12 | 4.12 | 3.11 | 3.05 | 2.67 | 2.43 |
| EMD-42774 | 8UXQ | 8.78 | 8.24 | 8.53 | 7.94 | 7.94 | 7.84 |
| EMD-42795 | 8UY8 | 3.69 | 3.69 | 3.06 | 3 | 2.24 | 2.12 |
| EMD-43346 | 8VLW | 4.26 | 4.24 | 3.21 | 3.46 | 2.4 | 2.26 |
| EMD-39454 | 8YON | 10.67 | 9.86 | 7.29 | 8.63 | 7.71 | 7.82 |
| EMD-44441 | 9BCX | 9.03 | 8.53 | 8.08 | 19.16 | 8.43 | 8.23 |
| EMD-45805 | 9CPO | 3.99 | 3.99 | 3.53 | 3.44 | 2.9 | 2.44 |
| EMD-46645 | 9D8P | 4.34 | 4.46 | 3.25 | 3.33 | 2.83 | 2.74 |
| EMD-62812 | 9L4F | 8.83 | 8.42 | 6.52 | 7.99 | 7.91 | 7.75 |

**Supplementary Table 9 | Summary of Q-score for the 27 half-maps processed by different methods.**

| Q-score | | | | | | | |
|---|---|---|---|---|---|---|---|
| EMDBID | PDBID | Averaged | phenix | DeepEMhancer | CryoTEN | EMReady | DEMO-EMReF |
| EMD-30535 | 7D0I | 0.600 | 0.628 | 0.571 | 0.598 | 0.670 | 0.669 |
| EMD-31595 | 7FIF | 0.241 | 0.342 | 0.250 | 0.315 | 0.330 | 0.316 |
| EMD-22375 | 7JLO | 0.576 | 0.634 | 0.581 | 0.619 | 0.676 | 0.674 |
| EMD-12296 | 7NF6 | 0.593 | 0.646 | 0.589 | 0.633 | 0.679 | 0.676 |
| EMD-13587 | 7PQ5 | 0.652 | 0.572 | 0.645 | 0.695 | 0.720 | 0.727 |
| EMD-26377 | 7U6O | 0.455 | 0.477 | 0.406 | 0.451 | 0.516 | 0.497 |
| EMD-33995 | 7YP7 | 0.539 | 0.594 | 0.536 | 0.584 | 0.635 | 0.618 |
| EMD-16555 | 8CC6 | 0.588 | 0.635 | 0.586 | 0.634 | 0.690 | 0.696 |
| EMD-27533 | 8DMB | 0.598 | 0.604 | 0.588 | 0.613 | 0.615 | 0.655 |
| EMD-29686 | 8G30 | 0.597 | 0.629 | 0.582 | 0.634 | 0.665 | 0.658 |
| EMD-29936 | 8GCN | 0.356 | 0.348 | 0.288 | 0.348 | 0.379 | 0.401 |
| EMD-35304 | 8IAJ | 0.606 | 0.668 | 0.621 | 0.668 | 0.713 | 0.710 |
| EMD-35912 | 8J12 | 0.587 | 0.588 | 0.021 | 0.578 | 0.620 | 0.635 |
| EMD-35948 | 8J2F | 0.550 | 0.634 | 0.570 | 0.609 | 0.672 | 0.671 |
| EMD-18345 | 8QE8 | 0.621 | 0.594 | 0.608 | 0.604 | 0.675 | 0.678 |
| EMD-18453 | 8QJX | 0.598 | 0.590 | 0.583 | 0.599 | 0.651 | 0.644 |
| EMD-18608 | 8QQX | 0.649 | 0.665 | 0.630 | 0.693 | 0.719 | 0.707 |
| EMD-18778 | 8QZM | 0.617 | 0.637 | 0.610 | 0.648 | 0.665 | 0.704 |
| EMD-41355 | 8TKO | 0.602 | 0.611 | 0.581 | 0.635 | 0.674 | 0.674 |
| EMD-42774 | 8UXQ | 0.164 | 0.152 | 0.119 | 0.168 | 0.191 | 0.163 |
| EMD-42795 | 8UY8 | 0.611 | 0.651 | 0.603 | 0.643 | 0.702 | 0.712 |
| EMD-43346 | 8VLW | 0.518 | 0.557 | 0.517 | 0.543 | 0.659 | 0.666 |
| EMD-39454 | 8YON | 0.125 | 0.169 | 0.078 | 0.114 | 0.112 | 0.110 |
| EMD-44441 | 9BCX | 0.119 | 0.166 | 0.074 | 0.110 | 0.146 | 0.139 |
| EMD-45805 | 9CPO | 0.540 | 0.577 | 0.525 | 0.567 | 0.611 | 0.602 |
| EMD-46645 | 9D8P | 0.643 | 0.636 | 0.623 | 0.645 | 0.693 | 0.693 |
| EMD-62812 | 9L4F | 0.234 | 0.342 | 0.254 | 0.291 | 0.283 | 0.310 |

**Supplementary Table 10 | Summary of CC_mask for the 27 half-maps processed by different methods.**

| CC_mask | | | | | | | |
|---|---|---|---|---|---|---|---|
| EMDBID | PDBID | Averaged | phenix | DeepEMhancer | CryoTEN | EMReady | DEMO-EMReF |
| EMD-30535 | 7D0I | 0.820 | 0.839 | 0.804 | 0.832 | 0.861 | 0.850 |
| EMD-31595 | 7FIF | 0.636 | 0.824 | 0.788 | 0.813 | 0.758 | 0.804 |
| EMD-22375 | 7JLO | 0.786 | 0.863 | 0.818 | 0.861 | 0.865 | 0.848 |
| EMD-12296 | 7NF6 | 0.846 | 0.856 | 0.823 | 0.879 | 0.878 | 0.871 |
| EMD-13587 | 7PQ5 | 0.825 | 0.748 | 0.836 | 0.836 | 0.894 | 0.895 |
| EMD-26377 | 7U6O | 0.778 | 0.635 | 0.732 | 0.847 | 0.789 | 0.817 |
| EMD-33995 | 7YP7 | 0.799 | 0.819 | 0.811 | 0.851 | 0.867 | 0.854 |
| EMD-16555 | 8CC6 | 0.828 | 0.871 | 0.836 | 0.874 | 0.880 | 0.876 |
| EMD-27533 | 8DMB | 0.801 | 0.832 | 0.818 | 0.849 | 0.853 | 0.830 |
| EMD-29686 | 8G30 | 0.871 | 0.810 | 0.840 | 0.855 | 0.869 | 0.885 |
| EMD-29936 | 8GCN | 0.821 | 0.485 | 0.724 | 0.695 | 0.745 | 0.822 |
| EMD-35304 | 8IAJ | 0.795 | 0.883 | 0.833 | 0.873 | 0.887 | 0.872 |
| EMD-35912 | 8J12 | 0.843 | 0.834 | 0.837 | 0.868 | 0.869 | 0.880 |
| EMD-35948 | 8J2F | 0.777 | 0.872 | 0.799 | 0.828 | 0.855 | 0.844 |
| EMD-18345 | 8QE8 | 0.792 | 0.857 | 0.804 | 0.781 | 0.858 | 0.858 |
| EMD-18453 | 8QJX | 0.859 | 0.860 | 0.844 | 0.896 | 0.878 | 0.881 |
| EMD-18608 | 8QQX | 0.871 | 0.906 | 0.853 | 0.866 | 0.878 | 0.866 |
| EMD-18778 | 8QZM | 0.875 | 0.879 | 0.854 | 0.842 | 0.876 | 0.877 |
| EMD-41355 | 8TKO | 0.836 | 0.803 | 0.773 | 0.793 | 0.817 | 0.841 |
| EMD-42774 | 8UXQ | 0.566 | 0.636 | 0.566 | 0.597 | 0.564 | 0.611 |
| EMD-42795 | 8UY8 | 0.865 | 0.874 | 0.856 | 0.820 | 0.886 | 0.885 |
| EMD-43346 | 8VLW | 0.830 | 0.762 | 0.832 | 0.765 | 0.855 | 0.858 |
| EMD-39454 | 8YON | 0.712 | 0.774 | 0.583 | 0.410 | 0.563 | 0.584 |
| EMD-44441 | 9BCX | 0.517 | 0.639 | 0.564 | 0.387 | 0.596 | 0.584 |
| EMD-45805 | 9CPO | 0.817 | 0.789 | 0.811 | 0.843 | 0.838 | 0.835 |
| EMD-46645 | 9D8P | 0.825 | 0.807 | 0.797 | 0.887 | 0.831 | 0.855 |
| EMD-62812 | 9L4F | 0.595 | 0.744 | 0.689 | 0.592 | 0.607 | 0.646 |

**Supplementary Table 11 | Summary of CC_box for the 27 half-maps processed by different methods.**

| CC_box | | | | | | | |
|---|---|---|---|---|---|---|---|
| EMDBID | PDBID | Averaged | phenix | DeepEMhancer | CryoTEN | EMReady | DEMO-EMReF |
| EMD-30535 | 7D0I | 0.688 | 0.714 | 0.800 | 0.882 | 0.895 | 0.889 |
| EMD-31595 | 7FIF | 0.468 | 0.618 | 0.796 | 0.902 | 0.878 | 0.888 |
| EMD-22375 | 7JLO | 0.685 | 0.748 | 0.850 | 0.907 | 0.911 | 0.901 |
| EMD-12296 | 7NF6 | 0.697 | 0.707 | 0.761 | 0.895 | 0.892 | 0.893 |
| EMD-13587 | 7PQ5 | 0.696 | 0.822 | 0.853 | 0.873 | 0.923 | 0.923 |
| EMD-26377 | 7U6O | 0.683 | 0.632 | 0.734 | 0.920 | 0.849 | 0.862 |
| EMD-33995 | 7YP7 | 0.651 | 0.691 | 0.818 | 0.909 | 0.917 | 0.920 |
| EMD-16555 | 8CC6 | 0.767 | 0.790 | 0.851 | 0.915 | 0.921 | 0.922 |
| EMD-27533 | 8DMB | 0.750 | 0.837 | 0.856 | 0.912 | 0.912 | 0.902 |
| EMD-29686 | 8G30 | 0.686 | 0.700 | 0.765 | 0.841 | 0.850 | 0.870 |
| EMD-29936 | 8GCN | 0.731 | 0.458 | 0.750 | 0.851 | 0.871 | 0.908 |
| EMD-35304 | 8IAJ | 0.732 | 0.785 | 0.868 | 0.914 | 0.926 | 0.919 |
| EMD-35912 | 8J12 | 0.782 | 0.803 | 0.842 | 0.937 | 0.918 | 0.925 |
| EMD-35948 | 8J2F | 0.656 | 0.718 | 0.789 | 0.881 | 0.916 | 0.920 |
| EMD-18345 | 8QE8 | 0.622 | 0.767 | 0.842 | 0.834 | 0.902 | 0.897 |
| EMD-18453 | 8QJX | 0.794 | 0.847 | 0.836 | 0.945 | 0.917 | 0.919 |
| EMD-18608 | 8QQX | 0.786 | 0.817 | 0.863 | 0.904 | 0.927 | 0.923 |
| EMD-18778 | 8QZM | 0.821 | 0.840 | 0.859 | 0.890 | 0.932 | 0.938 |
| EMD-41355 | 8TKO | 0.758 | 0.786 | 0.767 | 0.850 | 0.876 | 0.887 |
| EMD-42774 | 8UXQ | 0.713 | 0.707 | 0.652 | 0.743 | 0.725 | 0.765 |
| EMD-42795 | 8UY8 | 0.570 | 0.661 | 0.835 | 0.853 | 0.923 | 0.922 |
| EMD-43346 | 8VLW | 0.583 | 0.567 | 0.808 | 0.812 | 0.905 | 0.906 |
| EMD-39454 | 8YON | 0.850 | 0.882 | 0.722 | 0.661 | 0.750 | 0.783 |
| EMD-44441 | 9BCX | 0.692 | 0.738 | 0.656 | 0.559 | 0.774 | 0.791 |
| EMD-45805 | 9CPO | 0.775 | 0.703 | 0.834 | 0.902 | 0.899 | 0.899 |
| EMD-46645 | 9D8P | 0.660 | 0.728 | 0.755 | 0.882 | 0.843 | 0.838 |
| EMD-62812 | 9L4F | 0.626 | 0.762 | 0.776 | 0.796 | 0.809 | 0.823 |

**Supplementary Table 12 | Summary of CC_peaks for the 27 half-maps processed by different methods.**

| CC_peaks | | | | | | | |
|---|---|---|---|---|---|---|---|
| EMDBID | PDBID | Averaged | phenix | DeepEMhancer | CryoTEN | EMReady | DEMO-EMReF |
| EMD-30535 | 7D0I | 0.688 | 0.681 | 0.770 | 0.826 | 0.851 | 0.842 |
| EMD-31595 | 7FIF | 0.221 | 0.549 | 0.766 | 0.789 | 0.739 | 0.757 |
| EMD-22375 | 7JLO | 0.685 | 0.717 | 0.822 | 0.867 | 0.875 | 0.858 |
| EMD-12296 | 7NF6 | 0.697 | 0.667 | 0.730 | 0.827 | 0.825 | 0.824 |
| EMD-13587 | 7PQ5 | 0.696 | 0.748 | 0.837 | 0.830 | 0.884 | 0.888 |
| EMD-26377 | 7U6O | 0.683 | 0.512 | 0.708 | 0.826 | 0.775 | 0.809 |
| EMD-33995 | 7YP7 | 0.651 | 0.625 | 0.759 | 0.842 | 0.858 | 0.858 |
| EMD-16555 | 8CC6 | 0.767 | 0.768 | 0.822 | 0.875 | 0.885 | 0.884 |
| EMD-27533 | 8DMB | 0.750 | 0.799 | 0.814 | 0.854 | 0.860 | 0.834 |
| EMD-29686 | 8G30 | 0.686 | 0.615 | 0.670 | 0.708 | 0.709 | 0.752 |
| EMD-29936 | 8GCN | 0.731 | 0.304 | 0.649 | 0.644 | 0.703 | 0.778 |
| EMD-35304 | 8IAJ | 0.732 | 0.774 | 0.840 | 0.877 | 0.894 | 0.882 |
| EMD-35912 | 8J12 | 0.782 | 0.757 | 0.828 | 0.870 | 0.867 | 0.884 |
| EMD-35948 | 8J2F | 0.656 | 0.677 | 0.762 | 0.801 | 0.841 | 0.842 |
| EMD-18345 | 8QE8 | 0.622 | 0.767 | 0.807 | 0.769 | 0.863 | 0.853 |
| EMD-18453 | 8QJX | 0.794 | 0.782 | 0.810 | 0.895 | 0.868 | 0.878 |
| EMD-18608 | 8QQX | 0.786 | 0.789 | 0.828 | 0.858 | 0.880 | 0.873 |
| EMD-18778 | 8QZM | 0.821 | 0.776 | 0.826 | 0.812 | 0.864 | 0.860 |
| EMD-41355 | 8TKO | 0.758 | 0.707 | 0.742 | 0.787 | 0.814 | 0.840 |
| EMD-42774 | 8UXQ | 0.444 | 0.503 | 0.527 | 0.577 | 0.552 | 0.596 |
| EMD-42795 | 8UY8 | 0.570 | 0.616 | 0.825 | 0.796 | 0.876 | 0.874 |
| EMD-43346 | 8VLW | 0.583 | 0.508 | 0.809 | 0.745 | 0.847 | 0.847 |
| EMD-39454 | 8YON | 0.629 | 0.702 | 0.556 | 0.388 | 0.546 | 0.558 |
| EMD-44441 | 9BCX | 0.397 | 0.513 | 0.539 | 0.372 | 0.552 | 0.539 |
| EMD-45805 | 9CPO | 0.775 | 0.666 | 0.807 | 0.844 | 0.842 | 0.845 |
| EMD-46645 | 9D8P | 0.660 | 0.634 | 0.677 | 0.736 | 0.704 | 0.685 |
| EMD-62812 | 9L4F | 0.482 | 0.666 | 0.681 | 0.562 | 0.573 | 0.605 |

**Supplementary Table 13 | Summary of FSC-0.5 resolution for the 43 half-maps in the cross-validation experiment.** FSC-0.5 resolution was calculated between the DEMO-EMReF-processed half-map and its corresponding unprocessed complementary half-map.

| EMDBID | PDBID | EM/ET | halfmap1-halfmap2 | DEMO-EMReF-processed half-map1-halfmap2 | half-map1-DEMO-EMReF-processed halfmap2 |
|---|---|---|---|---|---|
| EMD-30535 | 7D0I | EM | 3.73 | 3.62 | 3.62 |
| EMD-22375 | 7JLO | EM | 4.27 | 4.04 | 4.03 |
| EMD-12296 | 7NF6 | EM | 3.99 | 3.86 | 3.85 |
| EMD-13587 | 7PQ5 | EM | 4.02 | 3.79 | 3.77 |
| EMD-26377 | 7U6O | EM | 3.98 | 3.93 | 3.94 |
| EMD-33995 | 7YP7 | EM | 7.5 | 5.88 | 4.55 |
| EMD-16555 | 8CC6 | EM | 4.02 | 3.89 | 3.92 |
| EMD-27533 | 8DMB | EM | 4.38 | 4.05 | 4.09 |
| EMD-29686 | 8G30 | EM | 4.3 | 4.07 | 4.06 |
| EMD-29936 | 8GCN | EM | 4.32 | 4.34 | 4.35 |
| EMD-18345 | 8QE8 | EM | 10.69 | 9.06 | 9.05 |
| EMD-35304 | 8IAJ | EM | 4.04 | 3.78 | 3.81 |
| EMD-35912 | 8J12 | EM | 4.09 | 3.81 | 3.81 |
| EMD-35948 | 8J2F | EM | 3.67 | 3.62 | 3.59 |
| EMD-18453 | 8QJX | EM | 4.04 | 3.86 | 3.9 |
| EMD-18608 | 8QQX | EM | 4.01 | 3.83 | 3.83 |
| EMD-18778 | 8QZM | EM | 4.03 | 3.76 | 3.79 |
| EMD-41355 | 8TKO | EM | 7.59 | 4.32 | 4.36 |
| EMD-42795 | 8UY8 | EM | 4.08 | 3.87 | 3.9 |
| EMD-43346 | 8VLW | EM | 7.07 | 4.41 | 4.45 |
| EMD-45805 | 9CPO | EM | 4.23 | 4.08 | 4.07 |
| EMD-46645 | 9D8P | EM | 8.03 | 6.88 | 6.88 |
| EMD-31595 | 7FIF | EM | 8.57 | 8 | 8.06 |
| EMD-42774 | 8UXQ | EM | 7.53 | 7.2 | 7.17 |
| EMD-39454 | 8YON | EM | 17.77 | 10.86 | 10.76 |
| EMD-44441 | 9BCX | EM | 8.57 | 8.3 | 8.26 |
| EMD-62812 | 9L4F | EM | 9.66 | 9.08 | 8.96 |
| EMD-11078 | 6Z5J | ET | 10.61 | 10.61 | 10.82 |
| EMD-13275 | 7PAK | ET | 10.8 | 9.87 | 9.91 |
| EMD-13991 | 7QIN | ET | 8.56 | 8.35 | 8.36 |
| EMD-32906 | 7WZ8 | ET | 8.68 | 9.06 | 9.08 |
| EMD-16183 | 8BQE | ET | 4.35 | 4.13 | 4.1 |
| EMD-16489 | 8C8O | ET | 4.21 | 4.11 | 4.12 |
| EMD-16511 | 8C9M | ET | 4.12 | 4.32 | 4.34 |
| EMD-16772 | 8CO6 | ET | 7.6 | 7.23 | 7.23 |
| EMD-19708 | 8S41 | ET | 13.26 | 10.27 | 10.28 |
| EMD-41485 | 8TPU | ET | 8.69 | 6.94 | 6.95 |

| | | | | | |
|---|---|---|---|---|---|
| EMD-39107 | 8YAX | ET | 7.56 | 7.28 | 7.27 |
| EMD-44990 | 9BX1 | ET | 9.96 | 10.69 | 10.73 |
| EMD-45591 | 9CHO | ET | 10.02 | 9.74 | 9.64 |
| EMD-19999 | 9EVD | ET | 9.56 | 8.88 | 8.92 |
| EMD-50210 | 9F62 | ET | 8.84 | 8.17 | 8.15 |
| EMD-50836 | 9FWV | ET | 5.98 | 5.79 | 5.81 |

118

**Supplementary Table 14 | Comparison of automated model building using deposited maps and DEMO-EMReF-processed maps.** Values represent the average residue coverage and sequence match percentage across 25 test cases. Bold values indicate the best performance for each metric.

| Methods | Residue_coverage(%) | Sequence_match(%) |
|---|---|---|
| Deposited | 59.4 | 38.9 |
| **DEMO-EMReF** | **73.6** | **63.3** |

**Supplementary Table 15 | Performance of automated model building using deposited maps and DEMO-EMReF-processed maps across 25 test cases.**

| EMDBID | PDBID | Residue coverage (%) | | SEQ MATCH(%) | |
|---|---|---|---|---|---|
| | | deposited | DEMO-EMReF | deposited | DEMO-EMReF |
| EMD-31561 | 7FER | 69.20 | 83.20 | 23.40 | 61.80 |
| EMD-22901 | 7KJX | 60.00 | 76.50 | 55.50 | 73.80 |
| EMD-24073 | 7MXL | 56.60 | 66.20 | 28.90 | 54.90 |
| EMD-12296 | 7NF6 | 65.40 | 87.90 | 63.30 | 74.60 |
| EMD-12309 | 7NG8 | 78.20 | 81.10 | 63.20 | 83.30 |
| EMD-12600 | 7NUR | 75.50 | 83.40 | 56.60 | 84.40 |
| EMD-9915 | 6K4M | 43.10 | 63.30 | 64.90 | 67.10 |
| EMD-12808 | 7OCI | 68.70 | 82.20 | 48.40 | 63.60 |
| EMD-12993 | 7ON1 | 81.60 | 83.40 | 48.40 | 48.30 |
| EMD-13035 | 7OQZ | 75.30 | 83.00 | 66.50 | 83.60 |
| EMD-13587 | 7PQ5 | 80.30 | 88.40 | 48.00 | 79.90 |
| EMD-24473 | 7RI4 | 38.60 | 68.50 | 40.20 | 60.20 |
| EMD-32536 | 7WIU | 68.40 | 79.50 | 27.60 | 65.60 |
| EMD-32845 | 7WV4 | 9.50 | 14.90 | 10.00 | 16.80 |
| EMD-33192 | 7XHA | 57.80 | 76.30 | 37.90 | 72.70 |
| EMD-33995 | 7YP7 | 77.90 | 83.70 | 45.80 | 57.90 |
| EMD-29936 | 8GCN | 53.90 | 58.40 | 7.30 | 11.20 |
| EMD-35912 | 8J12 | 55.30 | 67.90 | 56.20 | 72.90 |
| EMD-35948 | 8J2F | 77.60 | 90.10 | 68.10 | 92.20 |
| EMD-28534 | 8EPX | 26.9 | 58.1 | 74.40 | 85.00 |
| EMD-18638 | 8QSP | 25.70 | 36.80 | 5.20 | 43.80 |
| EMD-33666 | 7Y7I | 72.2 | 78 | 14.20 | 83.80 |
| EMD-41355 | 8TKO | 61.50 | 77.50 | 44.00 | 82.70 |
| EMD-46645 | 9D8P | 56.10 | 83.60 | 28.60 | 64.10 |
| EMD-47476 | 9E3A | 50.40 | 88.70 | 27.80 | 73.60 |

**Supplementary Table 16 | Evaluation of DEMO-EMReF on maps with artificially added noise.** Correlation coefficients (CC) between simulated maps and DEMO-EMReF-processed maps under different noise levels. Numbers in bold indicate the best performance for each metric.

| Methods | CC_mask | CC_box | CC_peaks |
|---|---|---|---|
| Simulated map | 0.828 | 0.885 | 0.834 |
| Simulated map+noise | 0.781 | 0.749 | 0.689 |
| **DEMO-EMReF_processed** | **0.811** | **0.890** | **0.814** |

**Supplementary Table 17 | Summary of ablation results.** Ablation experiments for DEMO-EMReF were conducted on the primary-map dataset. Numbers in bold indicate the best performance for each metric. Asterisks denote statistically significant differences compared with the baseline model, determined using a two-sided Wilcoxon signed-rank test, with '*' indicating $p < 0.05$, '**' indicating $p < 0.001$, and '***' indicating $p < 0.0001$. The results for DEMO-EMReF differ from those reported in Supplementary Table 1 because the average values were computed over 111 cases, excluding three cases for which FSC-0.5 could not be calculated.

| Training Methods | FSC-0.5 (Å) | Q-score | CC_mask | CC_box | CC_peaks |
|---|---|---|---|---|---|
| SCUnet | 4.14*** | 0.499*** | **0.825*** | **0.895***** | 0.773 |
| SCUnet++ | 4.13*** | 0.505*** | 0.829** | 0.893*** | 0.765*** |
| DEMO-EMReF (×3 depth) | 3.70*** | 0.551 | 0.822 | 0.880* | 0.766** |
| DEMO-EMReF | **3.53** | **0.552** | 0.823 | 0.881 | **0.783** |
